# Dual Preconditioned Mesenchymal Stem Cells-Derived Culture Conditioned Media Augment Immunomodulation and Drive Metabolic Reprogramming in Acute Graft-versus-Host Disease

**DOI:** 10.1101/2025.11.02.686182

**Authors:** Mohini Mendiratta, Meenakshi Mendiratta, Sandeep Rai, Ritu Gupta, Sameer Bakhshi, Vatsla Dadhwal, Deepam Pushpam, Prabhat Singh Malik, Raja Pramanik, Mukul Aggarwal, Aditya Kumar Gupta, Rishi Dhawan, Tulika Seth, Manoranjan Mahapatra, Baibaswata Nayak, Thoudam Debraj Singh, Sachin Kumar, Riyaz Ahmed Mir, Lakshay Malhotra, Sabyasachi Bandyopadhyay, GuruRao Hariprasad, Hridayesh Prakash, Sujata Mohanty, Ranjit Kumar Sahoo

**Affiliations:** Department of Medical Oncology, Dr. B. R. Ambedkar Institute Rotary Cancer Hospital, All India Institute of Medical Sciences, New Delhi, India-110029; Stem Cell Facility (DBT-Centre of Excellence for Stem Cell Research), All India Institute of Medical Sciences, New Delhi, India-110029; Laboratory Oncology Unit, Dr. B. R. Ambedkar Institute Rotary Cancer Hospital, All India Institute of Medical Sciences, New Delhi, India-110029; Department of Obstetrics and Gynecology, All India Institute of Medical Sciences, New Delhi, India-110029; Department of Hematology, All India Institute of Medical Sciences, New Delhi, India-110029; Department of Pediatric Oncology, All India Institute of Medical Sciences, New Delhi, India-110029; Department of Gastroenterology and Human Nutrition, All India Institute of Medical Sciences, New Delhi, India-110029; Department of Biochemistry, All India Institute of Medical Sciences, New Delhi, India-110029; Department of Biochemistry, Sri Venkateshwar College, University of Delhi, New Delhi-110021, India; Proteomics Sub-facility, Centralized Core Research Facility, All India Institute of Medical Sciences, New Delhi-110029, India; Department of Biophysics, All India Institute of Medical Sciences, New Delhi, India-110029; Amity Centre for Translational Research, Amity University, Sector 125, Noida, India 201313

**Keywords:** Acute Graft-versus-Host-Disease, Mesenchymal Stem Cells, Apoptosis, Hypoxia, Culture conditioned media, Immunomodulation, Metabolic shift, Antioxidant

## Abstract

**Background:** The toxicity associated with conventional conditioning regimens limits treatment outcomes in autoimmune disorders such as acute graft-versus-host disease (aGVHD). Developing effective adjunctive interventions to restore immune balance is therefore essential. This study investigated the immunomodulatory potential of mesenchymal stem cells (MSCs)-derived culture-conditioned media (CCM) from naïve and preconditioned MSCs-hypoxic (MSCs^HYP^), apoptotic (MSCs^APO^), and dual preconditioned (MSCs^HYP+APO^) in aGVHD.

**Methods:** Human MSCs isolated from bone marrow and Wharton’s Jelly were preconditioned under hypoxia (1% O₂), apoptosis (1 µM staurosporine, 24 h), or both. The immunoregulatory and antioxidant properties of their CCM were evaluated through T-cell proliferation assays, Treg induction, macrophage polarization, mitochondrial function, and T-cell bioenergetics. Comparative proteomic profiling of CCM from WJ-MSCs and WJ-MSCs^HYP+APO^ co-cultured with aGVHD patient-derived activated PBMNCs was performed using LC-MS/MS, alongside *in vivo* validation in a chemotherapy-induced aGVHD murine model.

**Results:** WJ-MSCs^HYP+APO^-CCM exhibited superior immunomodulatory efficacy, suppressing T-cell proliferation, promoting Treg and Th2/Th9 differentiation, and driving M2 macrophage polarization. It reduced mitochondrial ROS, enhanced mitochondrial polarization, and shifted T-cell metabolism from glycolysis toward oxidative phosphorylation. Proteomic analysis revealed modulation of IL-12, IL-17, and JAK–STAT pathways, along with regulation of complement, coagulation, and metabolic cascades. Interaction with immune cells further enhanced its antioxidant and tissue-reparative properties via extracellular matrix remodeling.

**Conclusion:** Dual preconditioning under hypoxia and apoptosis amplifies the immunomodulatory, antioxidant, and reparative efficacy of WJ-MSC-CCM, offering a potent non-cellular therapeutic strategy for aGVHD management.

**GRAPHICAL ABSTRACT:** *Immunomodulatory and immune metabolic reprogramming potential of naïve and preconditioned MSCs (MSCs, MSCs^HYP^, MSCs^APO^, MSCs^HYP+APO^) derived CCM in Acute GVHD (created using Biorender.com)*

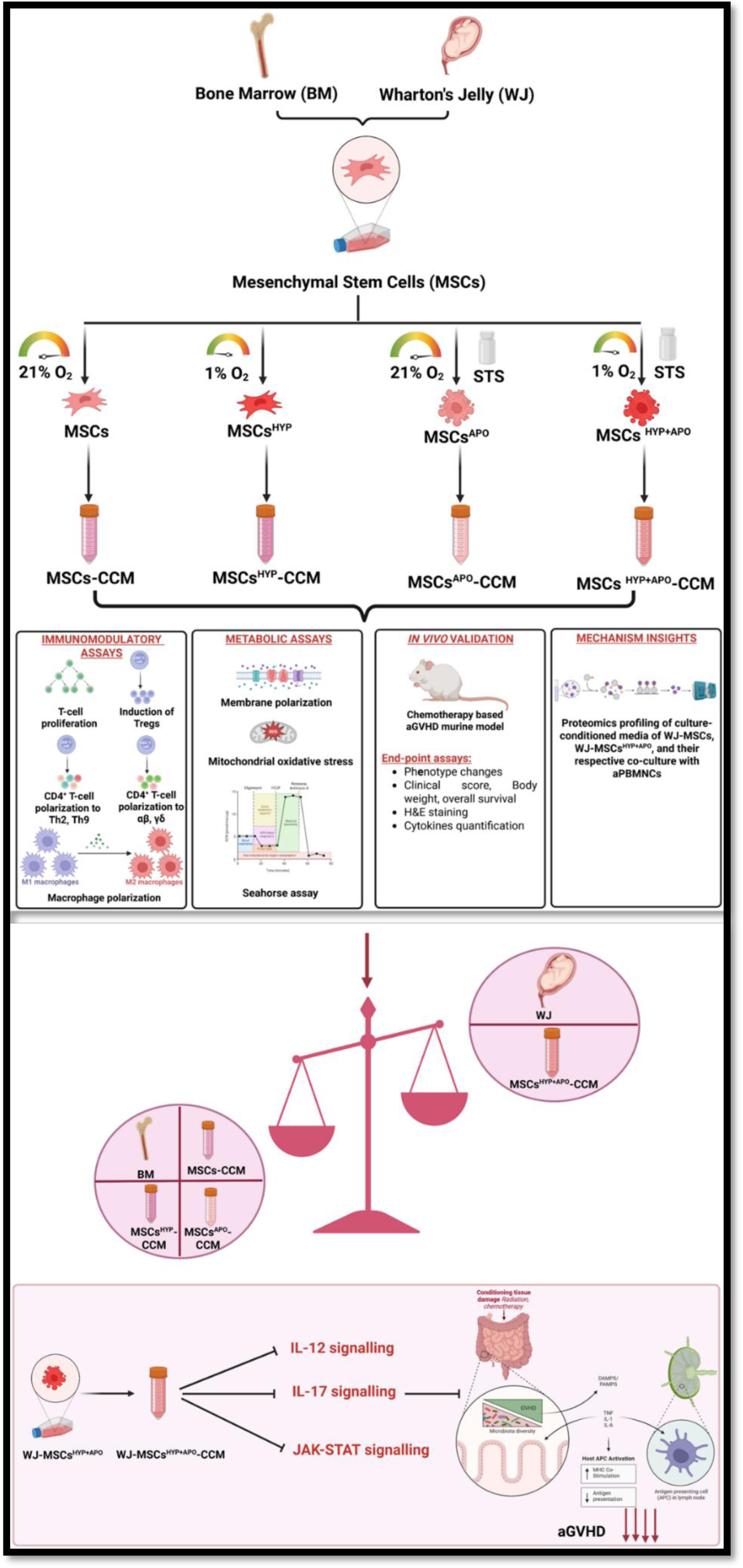

## INTRODUCTION

In recent years, mesenchymal stem cells (MSCs) have emerged as a promising immunomodulatory therapy due to their ability to regulate immune responses and promote tissue repair (1). However, viable MSCs-based therapies are associated with various challenges, such as donor variability, cell viability, and inconsistent outcomes, which have hindered their widespread application (2). Due to these limitations, there has been a shift toward exploring MSCs-derived secretome or culture-conditioned media (CCM) as a cell-free therapeutic approach, offering a more controlled and reproducible alternative for graft-versus-host-disease (GVHD) treatment. The therapeutic potential of MSCs depends mainly on their paracrine functions. An alternative or complement to cell therapy in inflammatory diseases is the use of media enriched with cytokines, chemokines, signaling molecules, and growth factors, and extracellular vesicles (EVs) secreted by *in vitro* cultures of MSCs, and their contents depend on their cellular origin and the physiological conditions in which they are produced (3).

Preconditioned and engineered MSCs can specifically improve immunomodulation effects in diverse clinical applications (4). Moreover, MSCs manipulations may cause cell functionality loss and genetic instability when performed outside their natural niches (5). Interestingly, numerous studies show that MSCs-derived EVs conditioned under hypoxia have a higher regenerative capacity than those obtained under normoxia. Hypoxia preconditioning of MSCs is being evaluated as a very attractive strategy for the isolation of EVs, with a high potential for clinical use (6).

Our previous study showed that both naïve and preconditioned MSCs, including those subjected to hypoxia or dual preconditioning (hypoxia combined with apoptosis), exhibited variable effects on T-cell proliferation *in vitro* during their direct coculture with T-cell derived from acute GVHD (aGVHD) patients, highlighting the complex and inconsistent immunomodulatory interactions, particularly given the central role of T-cell in aGVHD pathogenesis. To the best of our knowledge, this is the first study to systematically evaluate the comparative immunomodulatory effects of MSCs-derived CCM with different preconditioning strategies in aGVHD. Our findings provide critical insights into the therapeutic potential of the MSCs-derived CCM in modulating immune responses, alleviating oxidative stress, enhancing tissue repair and regeneration, and improving mitochondrial bioenergetics. These multifaceted effects highlight its promise as a cell-free therapeutic strategy for the effective management of aGVHD.

## RESULTS

### WJ-MSCs^HYP+APO^-CCM augmented T-cell polarization toward an anti-inflammatory and regulatory phenotype

To evaluate the immunomodulatory effects of MSCs-CCM, we analysed their impact on CD3^+^ T-cell proliferation and polarization using aGVHD patient-derived T-cell. Culture-conditioned media from both BM- and WJ-MSCs in naïve, hypoxia-preconditioned (^HYP^), apoptotic (^APO^), and combined hypoxia-apoptotic (^HYP+APO^) states were compared. A schematic representation outlining the assays performed with MSCs-CCM to assess their impact on T-cell (Figure S1). Across all conditions, MSCs-CCM significantly inhibited CD3^+^ T-cell proliferation, with WJ-MSCs demonstrating superior efficacy compared to BM-MSCs. Hypoxia preconditioning further enhanced this suppressive effect; BM-MSCs^HYP^-CCM reduced proliferation more effectively than BM-MSCs-CCM (46.60% vs 57.33%; p ≤ 0.0001), while WJ-MSCs^HYP^-CCM outperformed WJ-MSCs-CCM (39.19% vs 51.25%; p ≤ 0.0001). Notably, WJ-MSCs^HYP^-CCM was also superior to BM-MSCs^HYP^-CCM (p ≤ 0.0001). Similarly, both single (apoptotic) and dual (apoptotic and hypoxic) preconditioning significantly enhanced the immunosuppressive potential of MSCs. WJ-MSCs^HYP+APO^-CCM demonstrated the strongest inhibition of T-cell proliferation, significantly outperforming both its BM counterpart (21.03% vs 28.60%; p ≤ 0.0001) and WJ-MSC^APO^-CCM (21.03% vs 31.23%; p ≤ 0.0001) (Figure 1A).

**Figure 1:**
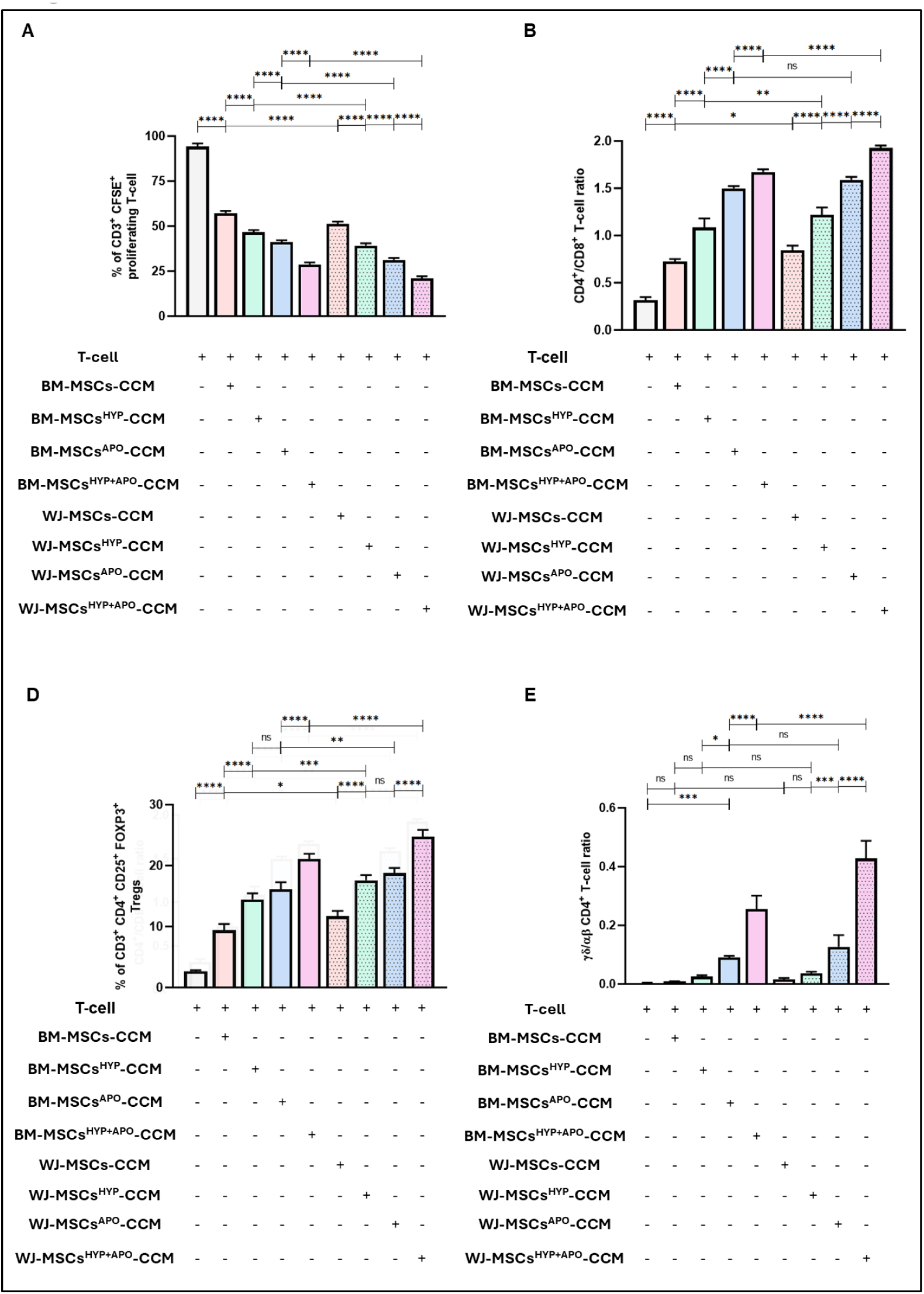
Effect of naïve, hypoxia, apoptotic, and hypoxia preconditioned apoptotic MSCs-CCM (MSCs, MSCs^HYP^, MSCs^APO^, MSCs^HYP+APO^) on the T-cell reprogramming derived from aGVHD patients. The bar graph represents (A) the percentage of CD3^+^ CFSE^+^ proliferating T-cell (n=25). (B) the ratio of CD4^+^/CD8^+^ T-cell (n=25). (C) the percentage of CD3^+^ CD4^+^ CD25^+^ FOXP3^+^ Tregs (n=25). (D) the ratio of γδ/αβ CD4^+^ T-cell (n=25) in the culture of MSCs-CCM and T-cell. Data shown represent the Mean±S.D of 25 independent experiments performed with T-cell derived from 25 different donors (biological replicates), with each experiment conducted in triplicate (technical replicates). Statistical analysis: Tukey’s multiple comparisons test; *≤0.05; **≤0.01; ***≤0.001; ****≤0.0001. Abbreviations: BM: Bone marrow; WJ: Wharton’s Jelly; MSCs: Mesenchymal Stem Cells; HYP: Hypoxia; APO: Apoptosis; CCM: Culture Conditioned Media

Moreover, the CD4^+^/CD8^+^ T-cell ratio revealed a consistent trend toward selective suppression of CD8^+^ T-cell. Hypoxia preconditioning significantly increased the CD4^+^/CD8^+^ ratio for both BM- and WJ-MSCs-CCM. WJ-MSCs^HYP^-CCM demonstrated higher efficacy than BM-MSCs^HYP^-CCM (1.22 vs 1.09; p ≤ 0.05), and this trend was more pronounced in the ^HYP+APO^ group, where WJ-MSCs^HYP+APO^-CCM showed a significantly higher ratio than BM-MSCs^HYP+APO^-CCM (1.93 vs 1.67; p ≤ 0.0001), indicating a robust shift toward CD4^+^ T-cell dominance (Figure 1B).

Further evaluation of CD4^+^ subsets revealed that MSCs-CCM promoted Treg induction, with a more pronounced effect observed under hypoxic and apoptotic conditions. BM-MSCs^HYP^-CCM induced significantly more Tregs compared to their naïve counterparts (14.48% vs 9.39%; p ≤ 0.0001), and WJ-MSCs^HYP^-CCM showed even greater enhancement (17.60% vs 11.68%; p ≤ 0.0001). In the apoptotic arms, WJ-MSCs^HYP+APO^-CCM demonstrated the highest Treg induction (24.73%), significantly greater than WJ-MSCs-CCM (18.77%; p ≤ 0.0001) and BM-MSCs^HYP+APO^-CCM (21.12%; p ≤ 0.0001) (Figure 1C).

In addition to Treg promotion, MSCs-CCM modulated γδ and αβ CD4^+^ T-cell subsets. Both BM- and WJ-MSCs^HYP^-CCM increased the γδ/αβ T-cell ratio significantly compared to their naïve states (BM: 0.025 vs 0.0093; WJ: 0.039 vs 0.016; both p ≤ 0.0001). This effect was further amplified in ^HYP+APO^ conditions. WJ-MSCs^HYP+APO^-CCM markedly increased the γδ/αβ ratio (0.43) compared to BM-MSCs^HYP+APO^-CCM (0.26; p ≤ 0.0001), indicating an enhanced ability to enrich regulatory γδ T-cell subsets under apoptotic-hypoxic preconditioning (Figure 1D).

MSCs-CCM also induced functional polarization of CD4^+^ T-cell toward anti-inflammatory phenotypes. The Th1/Th2 ratio significantly decreased with hypoxic preconditioning (BM-MSCs^HYP^-CCM vs BM-MSCs-CCM: 11.92 vs 14.95; p ≤ 0.01; WJ-MSCs^HYP^-CCM vs WJ-MSCs-CCM: 8.59 vs 12.84; p ≤ 0.001). The reduction was even more pronounced in the ^HYP+APO^ group, with WJ-MSCs^HYP+APO^-CCM reducing the Th1/Th2 ratio to 1.21 compared to 5.16 in WJ-MSCs^APO^-CCM (p ≤ 0.0001) (Figure S2A). A similar trend was observed in the Th1/Th17 ratio, where WJ-MSCs^HYP^-CCM outperformed BM-MSCs^HYP^-CCM (4.62 vs 6.34; p ≤ 0.0001), and WJ-MSCs^HYP+APO^-CCM demonstrated stronger Th17 suppression than BM-MSCs^HYP+APO^-CCM (7.85 vs 9.04; p ≤ 0.01) (Figure S2B).

Lastly, Th9 polarization was significantly influenced by both source and preconditioning. WJ-MSCs^HYP^-CCM demonstrated greater enhancement of Th9 cells compared to BM-MSCs^HYP^-CCM (13.42 vs 8.58; p ≤ 0.001). This effect was most prominent in the ^HYP+APO^ groups, where WJ-MSCs^HYP+APO^-CCM induced a significantly higher proportion of Th9 cells compared to BM-MSCs^HYP+APO^-CCM (8.22 vs 2.80; p ≤ 0.001), reinforcing the superior immunoregulatory capacity of WJ-MSCs^HYP+APO^-CCM (Figure S2C).

### WJ-MSCs^HYP+APO^-CCM enhanced polarization of M1 macrophages towards M2 anti-inflammatory phenotype

To evaluate the impact of MSCs-CCM, MSCs^HYP^-CCM, MSCs^APO^-CCM, and MSCs^HYP+APO^-CCM on macrophages derived from aGVHD patients (Figure S3), we assessed macrophage polarization under *in vitro* conditions using flow cytometry. Both MSCs-CCM and MSCs^HYP^-CCM showed a notable effect on suppressing the M1 phenotype, with a significant reduction observed between BM-MSCs^HYP^-CCM and BM-MSCs-CCM (42.89% vs. 53.80%; p ≤ 0.0001). A similar significant difference was observed between WJ-MSCs-CCM and WJ-MSCs^HYP^-CCM (47.73% vs. 32.42%; p ≤ 0.01). Notably, WJ-MSCs-CCM and WJ-MSCs^HYP^-CCM significantly outperformed their BM-MSCs counterparts in suppressing the M1 phenotype (47.73% vs. 53.80%, p ≤ 0.0001; 32.42% vs. 42.89%, p ≤ 0.0001). Similarly, significant inhibition of the M1 phenotype was observed between BM-MSCs^APO^-CCM and BM-MSCs^HYP+APO^-CCM (17.78% vs. 9.74%; p ≤ 0.0001), and between WJ-MSCs^APO^-CCM and WJ-MSCs^HYP+APO^-CCM (12.86% vs. 5.89%; p ≤ 0.0001). Once again, WJ-MSCs^APO^-CCM and WJ-MSCs^HYP+APO^-CCM significantly outperformed their BM-MSCs counterparts in suppressing the M1 phenotype (12.86% vs. 17.78%, p ≤ 0.0001; 5.89% vs. 9.74%, p ≤ 0.001) (Figure 2A).

**Figure 2:**
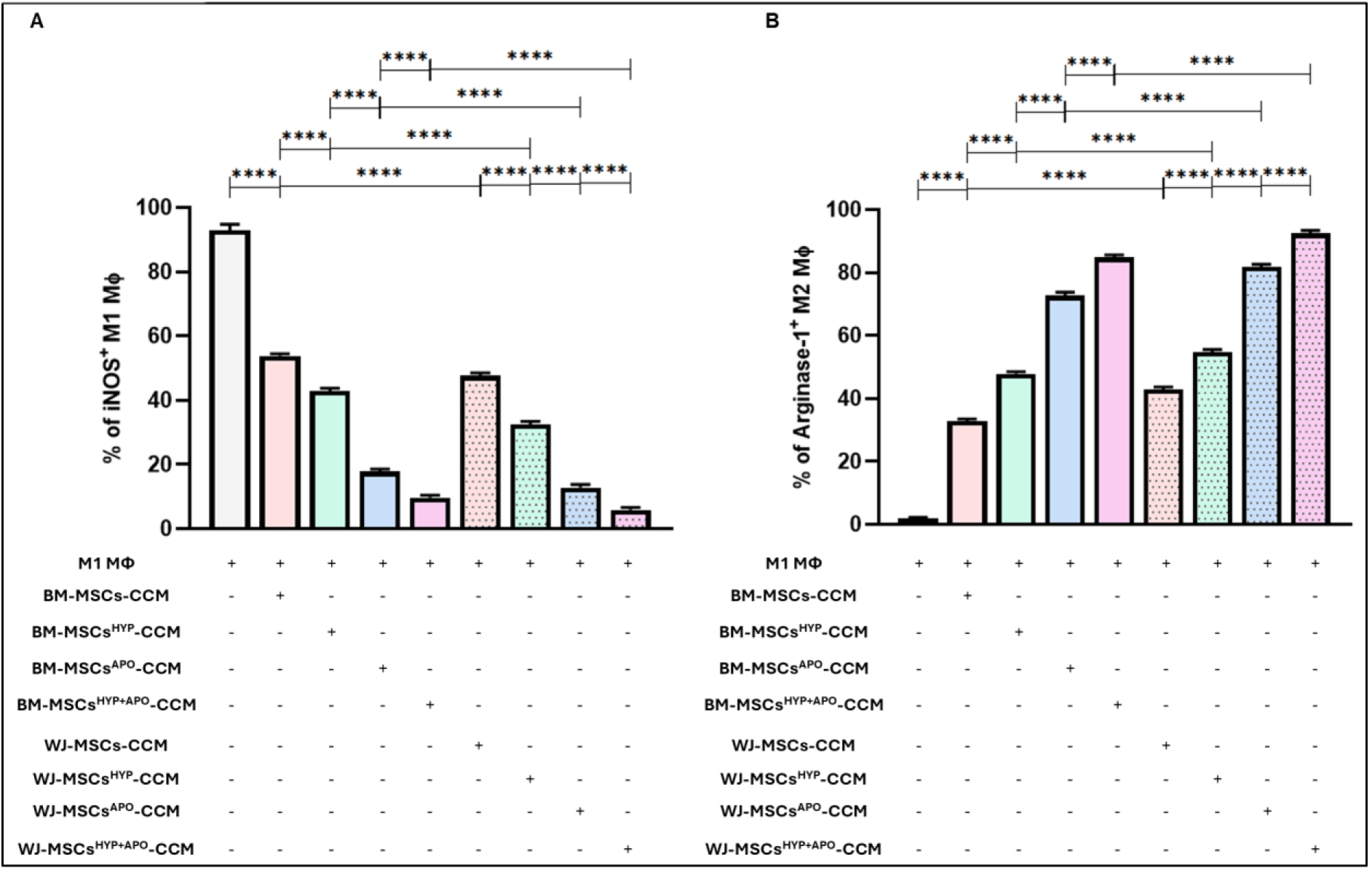
Effect of naïve, hypoxia, apoptotic, and hypoxia preconditioned apoptotic MSCs-CCM (MSCs, MSCs^HYP^, MSCs^APO^, MSCs^HYP+APO^) on the macrophage polarization of aGVHD patients. The bar graph represents (A) the percentage of iNOS^+^ M1 MФ. (B) the percentage of Arginase-1^+^ M2 MФ in the culture of MSCs-CCM and M1 MФ. Data shown represent the Mean±S.D of 25 independent experiments performed with M1 MФ derived from 25 different donors (biological replicates), with each experiment conducted in triplicate (technical replicates). Statistical analysis: Tukey’s multiple comparisons test; ***≤0.001; ****≤0.0001. Abbreviations: MФ: Macrophage; BM: Bone marrow; WJ: Wharton’s Jelly; MSCs: Mesenchymal Stem Cells; HYP: Hypoxia; APO: Apoptosis; CCM: Culture Conditioned Media

A corresponding trend was noted in macrophage polarization toward an M2 anti-inflammatory phenotype. WJ-MSCs^HYP^-CCM exhibited superior efficacy compared to BM-MSCs^HYP^-CCM, as indicated by a greater proportion of Arginase-1⁺ M2 macrophages (54.77% vs. 47.83%; p ≤ 0.0001). Likewise, WJ-MSCs^HYP+APO^-CCM demonstrated enhanced polarization towards the M2 phenotype compared to BM-MSCs^HYP+APO^-CCM (92.48% vs. 84.79%; p ≤ 0.0001) (Figure 2B).

### WJ-MSCs^HYP+APO^-CCM alleviated oxidative stress and modulated T-cell metabolism

To investigate the effect of MSCs-CCM on T-cell bioenergetics and their antioxidant potential, we performed in vitro assays by culturing T-cell derived from aGVHD patients with various types of MSCs-CCM (MSCs, MSCs^HYP^, MSCs^APO^, MSCs^HYP+APO^) (Figure S4). Mitochondrial ROS levels were initially assessed using MitoSOX dye. In line with the mitochondrial polarization data, WJ-MSCs-CCM demonstrated greater efficacy in reducing mitochondrial ROS levels compared to BM-MSCs-CCM (62.64% vs. 68.32%; p ≤ 0.01). Hypoxic preconditioning further enhanced this effect, although the difference between WJ-MSCs^HYP^-CCM and BM-MSCs^HYP^-CCM was not statistically significant (53.76% vs. 55.38%; p = 0.7314). Similarly, WJ-MSCs^APO^-CCM was more effective in reducing mitochondrial ROS than BM-MSCs^APO^-CCM (22.83% vs. 28.42%; p ≤ 0.001). A combination of hypoxia and apoptosis further enhanced ROS reduction, with WJ-MSCs^HYP+APO^-CCM demonstrating superior efficacy compared to BM-MSCs^HYP+APO^-CCM (12.64% vs. 18.25%; p ≤ 0.001) (Figure 3A).

**Figure 3:**
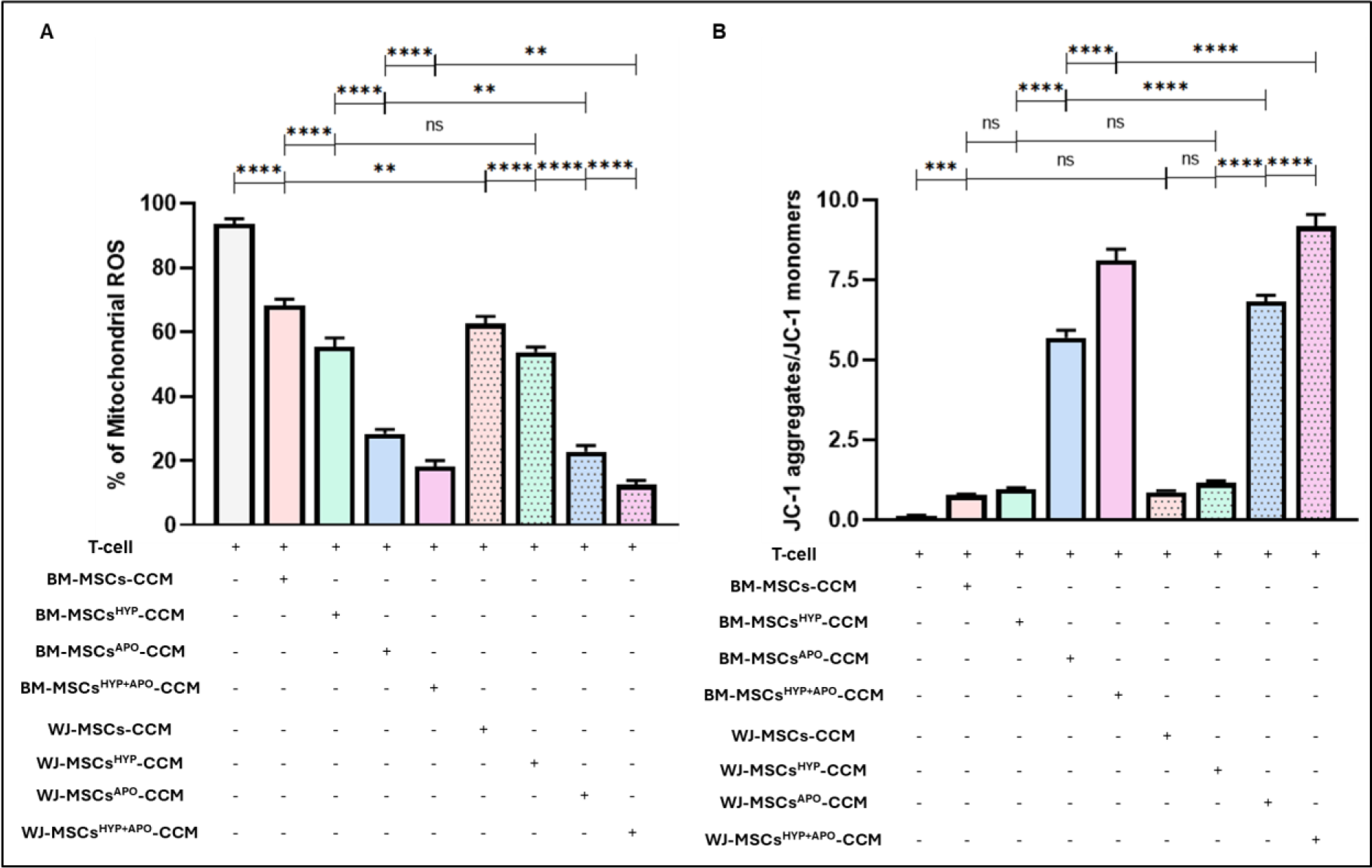
Effect of naïve, hypoxia, apoptotic, and hypoxia preconditioned apoptotic MSCs-CCM (MSCs, MSCs^HYP^, MSCs^APO^, MSCs^HYP+APO^) on the T-cell metabolism and oxidative stress of aGVHD patients. The bar graph represents (A) the percentage of mitochondrial ROS. (B) the ratio of JC-1 aggregates/JC-1 monomers in the culture of aGVHD patients-derived T-cell with MSCs-CCM. Data shown represent the Mean±S.D of 25 independent experiments performed with T-cell derived from 25 different donors (biological replicates), with each experiment conducted in triplicate (technical replicates). Statistical analysis: Tukey’s multiple comparisons test; *≤0.05; **≤0.01; ***≤0.001; ****≤0.0001. Abbreviations: BM: Bone marrow; WJ: Wharton’s Jelly; MSCs: Mesenchymal Stem Cells; HYP: Hypoxia; APO: Apoptosis; JC-1: 5,5′,6,6′-tetrachloro-1,1′,3,3′-tetraethylbenzimidazolocarbocyanine iodide; CCM: Culture-Conditioned Media

We further assessed mitochondrial health in T-cell from aGVHD patients using JC-1 dye to evaluate mitochondrial membrane potential. T-cell cultured under basal conditions exhibited mitochondrial depolarization, indicated by a reduced JC-1 aggregate/monomer ratio. Treatment with both MSCs-CCM and MSCs^HYP^-CCM improved mitochondrial polarization, with WJ-MSCs-CCM showing a significantly higher JC-1 ratio compared to BM-MSCs-CCM (0.87 vs. 0.77; p ≤ 0.01), and a similar trend was observed for MSCs^HYP^-CCM (1.16 vs. 0.97; p ≤ 0.0001). Likewise, MSCs^APO^-CCM and MSCs^HYP+APO^-CCM markedly restored mitochondrial polarization, with WJ-MSCs^APO^-CCM showing a significantly higher JC-1 ratio compared to BM-MSCs^APO^-CCM (6.82 vs. 5.70; p ≤ 0.0001), and a similar effect was observed for WJ-MSCs^HYP+APO^-CCM over BM-MSCs^HYP+APO^-CCM (9.17 vs. 8.11; p ≤ 0.0001) (Figure 3B).

To assess the influence of differentially preconditioned MSC-derived CCMs on T-cell mitochondrial function, we performed the Seahorse XF Mitochondrial Stress Test on CD3⁺ T-cell exposed to conditioned media obtained from naïve MSCs, hypoxia-preconditioned MSCs (^HYP^), apoptotic MSCs (^APO^), and dual preconditioned MSCs (^HYP+APO^), derived from both WJ and BM sources. The analysis revealed that all MSCs-CCM groups significantly enhanced mitochondrial respiration in T-cell compared to untreated controls, as reflected by improvements in basal respiration, maximal respiration, ATP production, SRC, and OCR, while simultaneously reducing proton leak and EACR, indicating a shift toward oxidative phosphorylation.

Among all tested groups, the WJ-MSCs^HYP+APO^-CCM induced the most pronounced improvement in mitochondrial parameters, outperforming other conditions across all variables-basal respiration, ATP production, proton leak, maximal respiration rate, SRC, ECAR, and OCR. MSCs^HYP^-CCM, MSCs^APO^-CCM also demonstrated enhanced mitochondrial support compared to naïve MSCs-CCM, though to a lesser extent than the dual preconditioned condition. These results indicate that combined hypoxic and apoptotic preconditioning synergistically enhances the capacity of MSCs to restore mitochondrial health in activated T-cell, promoting a metabolic reprogramming favoring OXPHOS over glycolysis, potentially underpinning the enhanced immunosuppressive effects observed in the aGVHD setting (Figure 4A-G).

**Figure 4:**
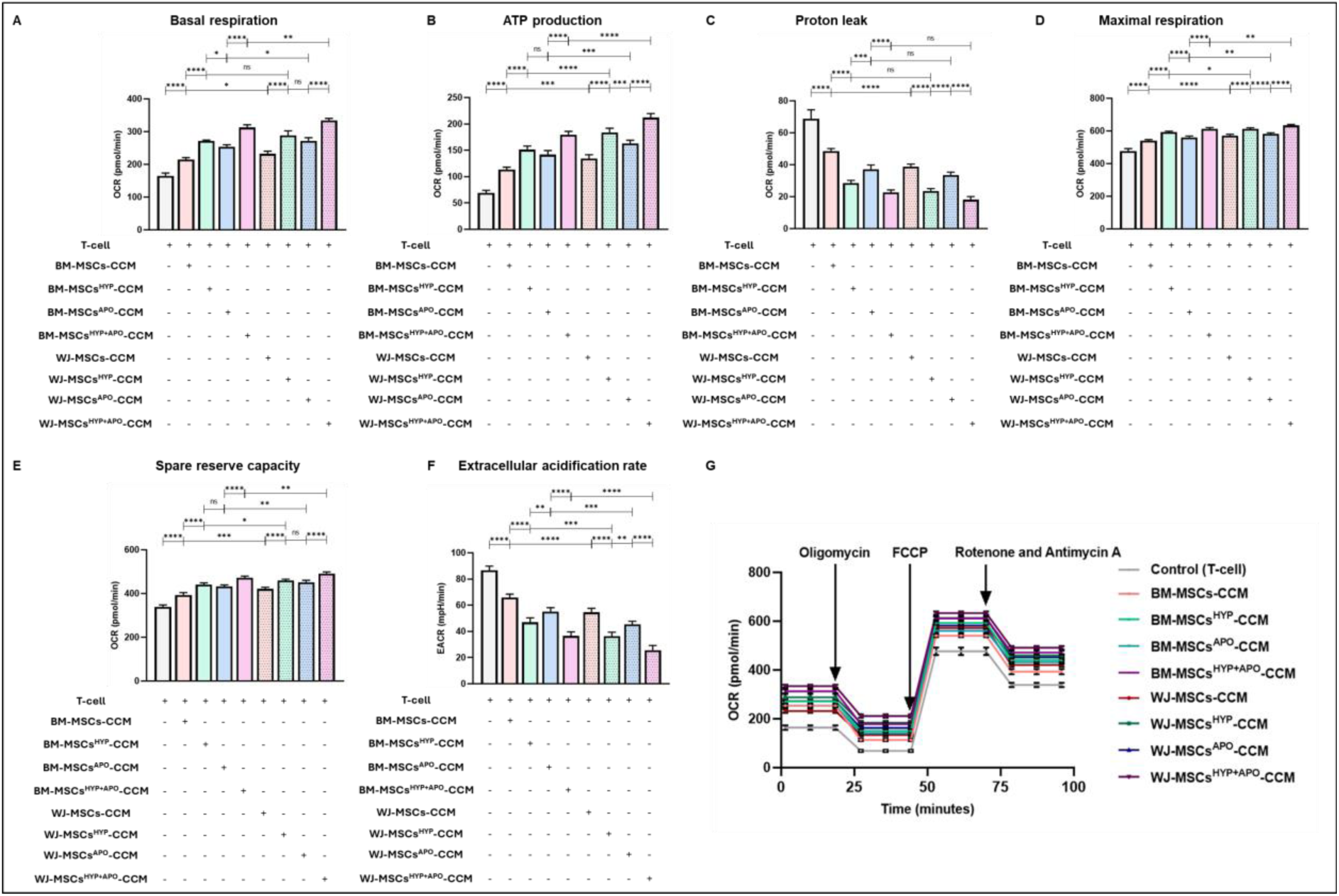
Effect of naïve, hypoxia, apoptotic, and hypoxia preconditioned apoptotic MSCs-CCM (MSCs, MSCs^HYP^, MSCs^APO^, MSCs^HYP+APO^) on the T-cell mitochondrial bioenergetics of aGVHD patients. The bar graph represents (A) Basal mitochondrial respiration. (B) ATP production. (C) Proton leak. (D) Maximal respiration. (E) Spare reserve capacity. (F) Extracellular acidification rate. (G) Real-time changes in the oxygen consumption rate in the culture of aGVHD patients-derived T-cell with MSCs-CCM. Data shown represent the Mean±S.D of 25 independent experiments performed with T-cell derived from 25 different donors (biological replicates), with each experiment conducted in triplicate (technical replicates). Statistical analysis: Tukey’s multiple comparisons test; *≤0.05; **≤0.01; ***≤0.001; ****≤0.0001. Abbreviations: BM: Bone marrow; WJ: Wharton’s Jelly; MSCs: Mesenchymal Stem Cells; HYP: Hypoxia; APO: Apoptosis; OCR: Oxygen consumption rate; EACR: extracellular acidification rate; CCM: Culture-Conditioned Media

### WJ-MSCs^HYP+APO^-CCM exhibited superior cytotoxic and pro-apoptotic effects on leukemic cell lines

To evaluate the anti-leukemic potential of MSCs-CCM, we assessed the cytotoxic effects of CCM obtained from both BM-MSCs and WJ-MSCs, including their naive and dual preconditioned (^HYP+APO^) variants, on two human leukemic cell lines-Kasumi-1 and MOLM-13. Cell viability was initially assessed using the MTS assay, followed by quantification of apoptotic, viable, and necrotic cell populations via Annexin V/7-AAD staining and flow cytometry. Overall, MSC-CCM exhibited significant cytotoxic effects in both cell lines, with dual-preconditioned MSC-CCM (BM-MSCs^HYP+APO^-CCM and WJ-MSCs^HYP+APO^-CCM) demonstrating markedly enhanced cytotoxicity compared to their respective naïve counterparts, irrespective of tissue source. When comparing the source of MSCs, WJ-MSCs^HYP+APO^-CCM consistently outperformed BM-MSCs^HYP+APO^-CCM. In the MTS assay, WJ-MSCs^HYP+APO^-CCM significantly reduced cell viability in both Kasumi-1 (83.08% vs. 52.84%; p ≤ 0.0001) and MOLM-13 (76.47% vs. 34.06%; p ≤ 0.0001) cell lines (Figure S5A-B). A similar trend was observed in the apoptosis assay, where WJ-MSCs^HYP+APO^-CCM induced a higher percentage of apoptotic cells in Kasumi-1 (53.73% vs. 43.38%; p ≤ 0.0001) and MOLM-13 (68.33% vs. 48.62%; p ≤ 0.0001) compared to BM-MSCs^HYP+APO^-CCM (Figure S5C-D). Furthermore, comparative analysis of the two leukemic cell lines revealed that Kasumi-1 cells were more resistant than MOLM-13, as reflected by a lower reduction in cell viability and a reduced proportion of apoptotic cells across all treatment groups.

### WJ-MSCs ^HYP+APO^-CCM augmented endothelial repair and tube formation

To evaluate the tissue repair potential of MSCs-CCM (WJ-MSCs-CCM and WJ-MSCs^HYP+APO^-CCM), H₂O₂-treated HUVECs were subjected to tube formation assays. Both CCMs promoted angiogenesis, as evident from tube-like structures in morphological images (Figure S6A); however, WJ-MSCs^HYP+APO^-CCM exhibited superior angiogenic potential. Quantitative analysis revealed significantly higher angiogenic parameters in the WJ-MSCs^HYP+APO^-CCM group compared to WJ-MSCs-CCM, including total number of loops (19 vs. 15; p ≤ 0.01), junctions (91 vs. 81; p ≤ 0.0001), branches (178.2 vs. 158.4; p ≤ 0.0001), and total branching length (29,750 mm vs. 19,500 mm; p ≤ 0.0001) (Figure S6B-E).

### WJ-MSCs^HYP+APO^-CCM augmented the alleviation of aGVHD in chemotherapy-based murine models

Our *in vitro* findings demonstrated that WJ-MSCs^HYP+APO^-CCM exhibited superior immunomodulatory effects compared to other groups, particularly in modulating T-cell responses, macrophage polarization toward an anti-inflammatory and regulatory phenotype, and influencing T-cell metabolism. These *in vitro* results were further validated using preclinical *in vivo* chemotherapy-based aGVHD murine models, with treatment groups comprising WJ-MSCs, WJ-MSCs^HYP+APO^, and their respective CCM.

We evaluated the therapeutic effects of WJ-MSCs, WJ-MSCs^HYP+APO^, and their CCM in a chemotherapy-based aGVHD murine model, which was designed to more closely mimic real-world transplant settings, depicted in Figure 5A. By days 4-5 post-transplantation (D+4-D+5), mice developed aGVHD, characterized by significant phenotypic changes, including hunching, skin ruffling, skin denuding, oral lesions, partial or complete eyelid closure, and, in some cases, forelimb disappearance, marking the onset of aGvHD (Figure 5B).

**Figure 5:**
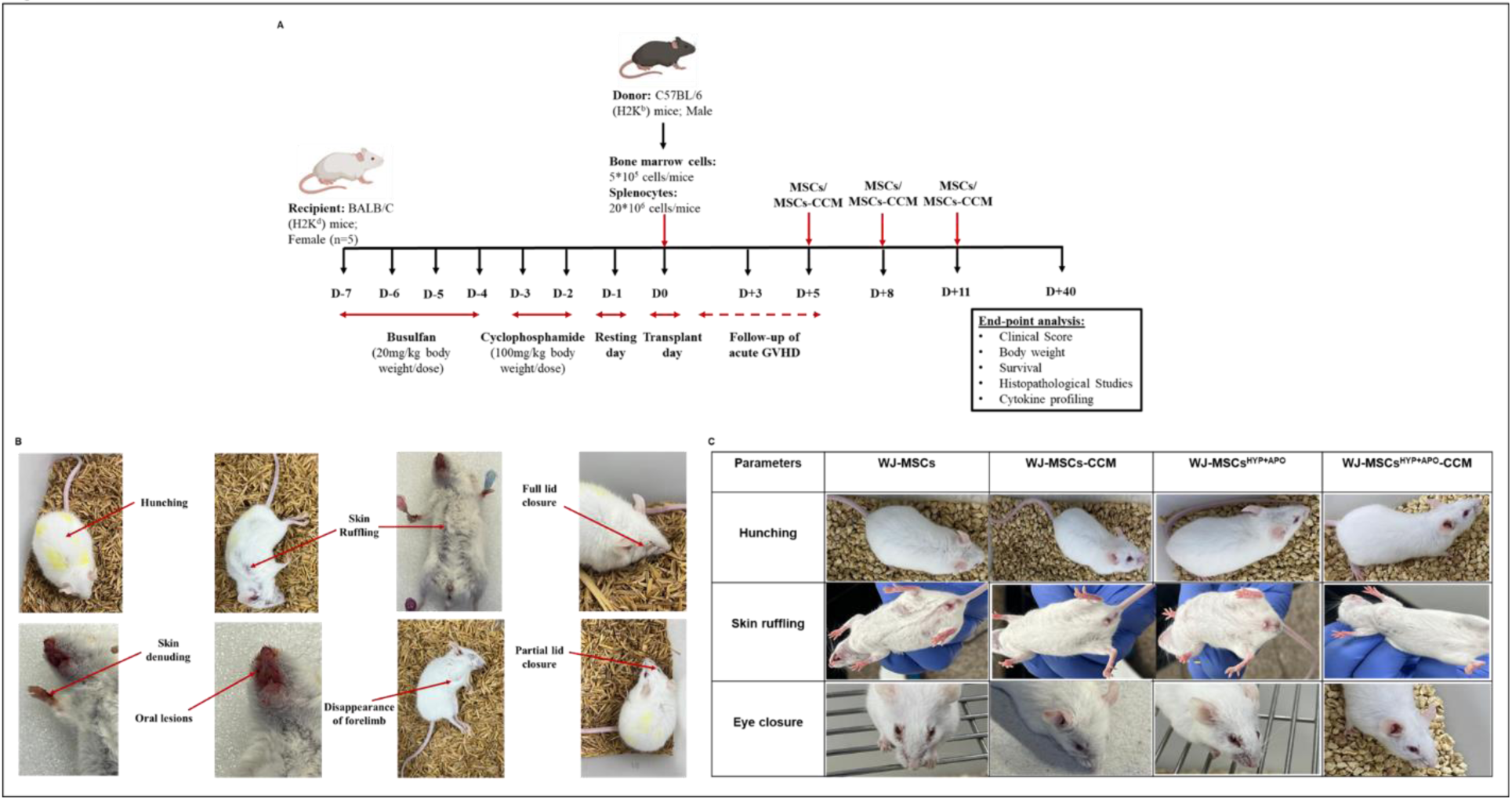
Effect of WJ-MSCs, WJ-MSCs^HYP+APO^, and their CCM in a chemotherapy-based aGVHD murine model. (A) Schematic representation of the development of chemotherapy-based aGVHD model and the timeline of administration of therapeutic intervention. (B) phenotype of the aGVHD control group. (C) phenotype in treated groups. Abbreviations: WJ: Wharton’s Jelly; MSCs: Mesenchymal Stem Cells; HYP: Hypoxia; APO: Apoptosis; CCM: Culture-Conditioned Media

Following therapeutic intervention, a notable improvement in phenotypic manifestations was observed, with a reduction in hunching, skin ruffling, and eyelid closure. These improvements were most pronounced in the WJ-MSCs^HYP+APO^-CCM-treated group, followed by WJ-MSCs^HYP+APO^-CCM, WJ-MSCs-CCM, and WJ-MSCs (Figure 5C).

We further analyzed the engraftment of donor cells in aGVHD control and treated groups and observed the complete presence of H-2Kb^+^ cells in the recipient mice, indicating that MSCs and their CCM did not affect donor cell engraftment (Figure S7). The overall survival was improved in all treated groups relative to the aGVHD control group (Figure S8A). Notably, body weight increased from D+12 to D+40 in the treated groups compared to the aGVHD control group (11.79 g vs 28.6 g) (Figure S8B). Additionally, A significant reduction in clinical scores was observed in all treated groups from D+12 to D+40 post-transplantation compared to the aGVHD control group (0.66 vs 10) (Figure S8C). Among the treatment groups, WJ-MSCs^HYP+APO^-CCM demonstrated the most pronounced benefits, exhibiting superior efficacy in reducing clinical scores, increasing body weight, and enhancing overall survival. The treated groups exhibited a significant shift in the cytokine profile compared to the control group, as shown in Figure S8D-F. This was characterized by a concurrent decrease in pro-inflammatory cytokines IFN-γ and IL-17α, alongside an increase in the anti-inflammatory cytokine IL-10. The reduction in IFN-γ, a key mediator of Th1-driven inflammation, suggests suppression of hyperactive immune responses that contribute to aGVHD pathology. Similarly, the decrease in IL-17α, associated with Th17 cells, indicates a downregulation of pathways linked to tissue damage and inflammation. Conversely, the elevated levels of IL-10 highlight enhanced anti-inflammatory signaling, promoting immune regulation and tissue repair. These changes in cytokine levels reflect an overall shift toward an immunomodulatory and tolerogenic environment in the treated groups, contributing to the observed amelioration of aGVHD symptoms and organ protection. These results also revealed that WJ-MSCs^HYP+APO^-CCM was better than other groups in alleviating aGVHD.

Moreover, histopathological analysis revealed significant protection of multiple organs, including the liver, lungs, intestine, and skin, from damage in the treated groups compared to the aGVHD control group, as shown in Figure 6A-E. In the liver, there was a marked reduction in lymphocytic infiltration (Figure 6F), preservation of hepatic architecture, and decreased periportal and parenchymal inflammation. The lungs exhibited reduced perivascular and alveolar lymphocyte infiltration (Figure 6G), with less alveolar wall thickening and minimal interstitial fibrosis. In the intestine, decreased lymphocytic infiltration (Figure 6H) was observed alongside improved villous architecture, reduced crypt destruction, and diminished inflammatory cell presence in the lamina propria. Similarly, in the skin, a reduction in dermal lymphocyte infiltration (Figure 6I) and improved epidermal integrity were evident, with fewer signs of dermo-epidermal separation or necrosis. These findings collectively indicate that the treatment provided substantial organ protection by mitigating immune-mediated damage and preserving tissue integrity.

**Figure 6:**
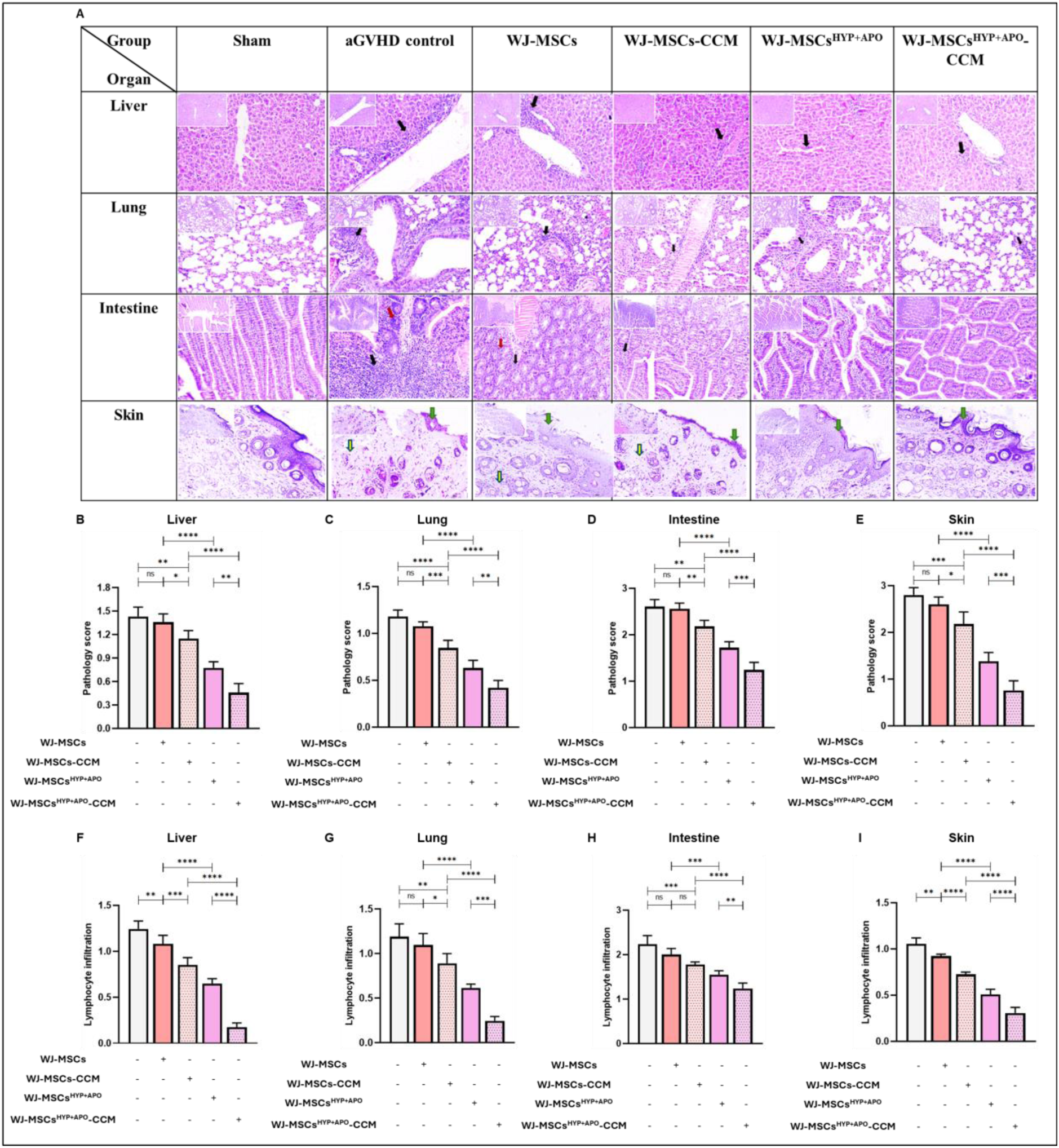
Effect of WJ-MSCs, WJ-MSCs^HYP+APO^, and their CCM on organ protection in a chemotherapy-based aGVHD murine model. (A) H&E staining of liver, lung, intestine, and skin tissue sections of sham, aGVHD control, WJ-MSCs, WJ-MSCs-CCM, WJ-MSCs^HYP+APO^, WJ-MSCs^HYP+APO^-CCM. The bar graphs represent the pathology score of (B) liver. (C) lung. (D) intestine. (E) skin tissue sections of the control and treated groups. Further, the bar graphs represent the lymphocyte infiltration in (F) liver. (G) lung. (H) intestine. (I) skin tissue sections of the control and treated groups. Data presented as Mean ±S.D. (N= 8 mice/group). This analysis was performed at D+21 in all treated groups and at D+7 in the aGVHD control group. Abbreviations: WJ: Wharton’s Jelly; MSCs: Mesenchymal Stem Cells; HYP: Hypoxia; APO: Apoptosis; CCM: Culture-Conditioned Media

### Immune cell interaction with WJ-MSCs^HYP+APO^ altered their secretome composition and mediated immunomodulation through metabolic reprogramming and enriched immune regulatory proteins

In our preclinical *in vitro* and *in vivo* studies, we observed that WJ-MSCs^HYP+APO^-CCM exhibited superior capabilities in immune restoration compared to others. To further elucidate the underlying mechanisms contributing to this enhanced performance, we conducted label-free proteomics analysis on the CCM of WJ-MSCs^HYP+APO^ (n=3) alone, as well as the CCM of the co-culture of WJ-MSCs^HYP+APO^ with aGVHD patients derived aPBMNCs (n=3), alongside separate analyses for WJ-MSCs-CCM (n=3) and CCM of the co-culture of WJ-MSCs with aGVHD patients derived aPBMNCs (n=3), depicted in Figure S9.

### WJ-MSCs-CCM versus WJ-MSCs^HYP+APO^-CCM

The comparative analysis of WJ-MSCs-CCM compared to WJ-MSCs^HYP+APO^-CCM revealed 323 total proteins. Among these, 80 proteins displayed statistically significant alterations with a Log_2_ fold change threshold of ≥ 1.5 and ≤ -1.5. Moreover, we observed that 43 dysregulated proteins were downregulated, while 37 were upregulated in the WJ-MSCs-CCM (Figure 7A-D).

**Figure 7:**
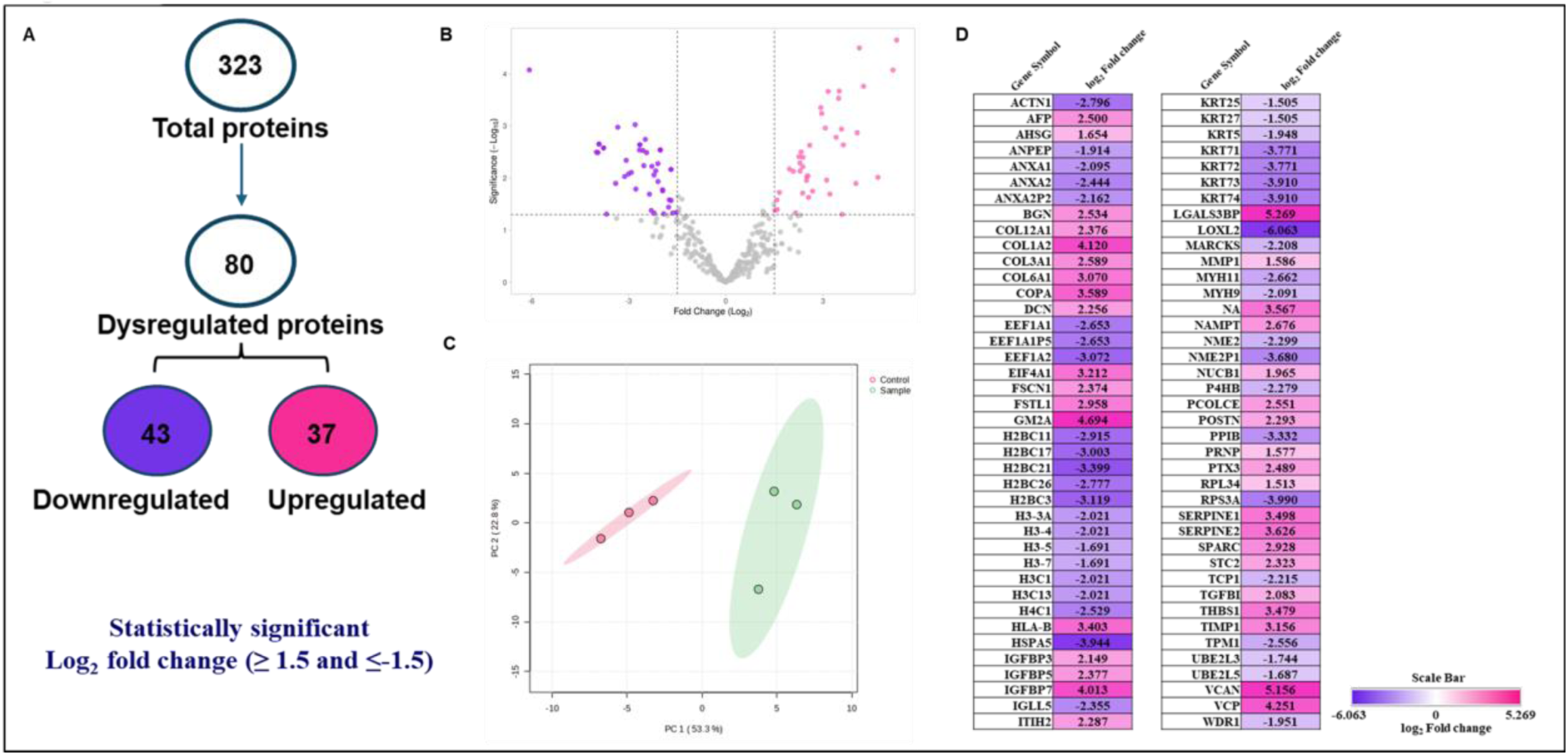
Label-free proteomics analysis of the CCM of WJ-MSCs and WJ-MSCs^HYP+APO^ using LC-MS/MS. (A) A flow chart depicts the total number of identified and dysregulated proteins. (B) A volcano plot highlights differentially expressed proteins. (C) Principal component analysis (PCA) demonstrates the good reproducibility of each biological replicate. (D) A heat map shows the expression levels of dysregulated proteins. Independent experiments were conducted with three different donors (biological replicates).

Biological and molecular function analyses of the upregulated proteins revealed that WJ-MSCs-CCM mediates immune regulation through multiple mechanisms, including negative regulation of protein processing (64.71%), cellular response to amino acid stimulus (11.76%), and peptide cross-linking (5.88%). Additionally, 71.34% of the enriched proteins were associated with growth factor binding, with contributions from the ECM structural component (14.29%), and collagen binding (14.29%) (Figure S10A-B). Similarly, cellular component analysis of upregulated proteins revealed that a significant proportion of the proteins, accounting for 50.0%, were localized within ER lumen and collagen-containing ECM. (Figure S10C).

Further functional enrichment analysis of these upregulated proteins highlighted a significant enrichment of proteins involved in the ECM proteoglycans (45.71%) and ECM organization (42.86%) (Figure S11A-B). KEGG pathway analysis underscored the role of the p53 signaling pathway in immune regulation (Figure S11C).

Interestingly, biological processes of down-regulated proteins demonstrated statistically significant enrichment, with intermediate filament organization (27.78%), phospholipase A2 inhibitor activity (16.67%), and nucleosome assembly (16.67%) (Figure S12A). Moreover, the analysis showcased various molecular functions, notably significant downregulation of phospholipase A2 inhibitor activity (83.33%) in WJ-MSCs-CCM compared to WJ-MSCs^HYP+APO^-CCM (Figure S12B), which are critical for immunomodulation.

The functional enrichment analysis of downregulated proteins in the WJ-MSCs-CCM revealed significant insights into immunomodulation. Notably, 66.67% of the identified proteins were associated with neutrophil extracellular trap formation, and 33.33% of the proteins were associated with systemic lupus erythematosus (SLE) (Figure S12C). Furthermore, the Reactome pathway analysis indicated that a significant proportion of proteins were involved in the activation of RHO GTPases, which activate PKNs (94.81%), involved in cell growth and differentiation (Figure S12D).

### WJ-MSCs-CCM versus CCM of coculture of WJ-MSCs with aGVHD patients-derived aPBMNCs

Next, the comparative analysis of WJ-MSCs-CCM compared to CCM of coculture of WJ-MSCs and aGVHD patients-derived aPBMNCs revealed 336 total proteins. Among these, 152 proteins displayed statistically significant alterations with a Log_2_ fold change threshold of ≥ 1.5 and ≤ -1.5. Moreover, we observed that 65 dysregulated proteins were downregulated, while 87 were upregulated in the co-culture of WJ-MSCs-CCM with aPBMNCs (Figure 8A-D).

**Figure 8:**
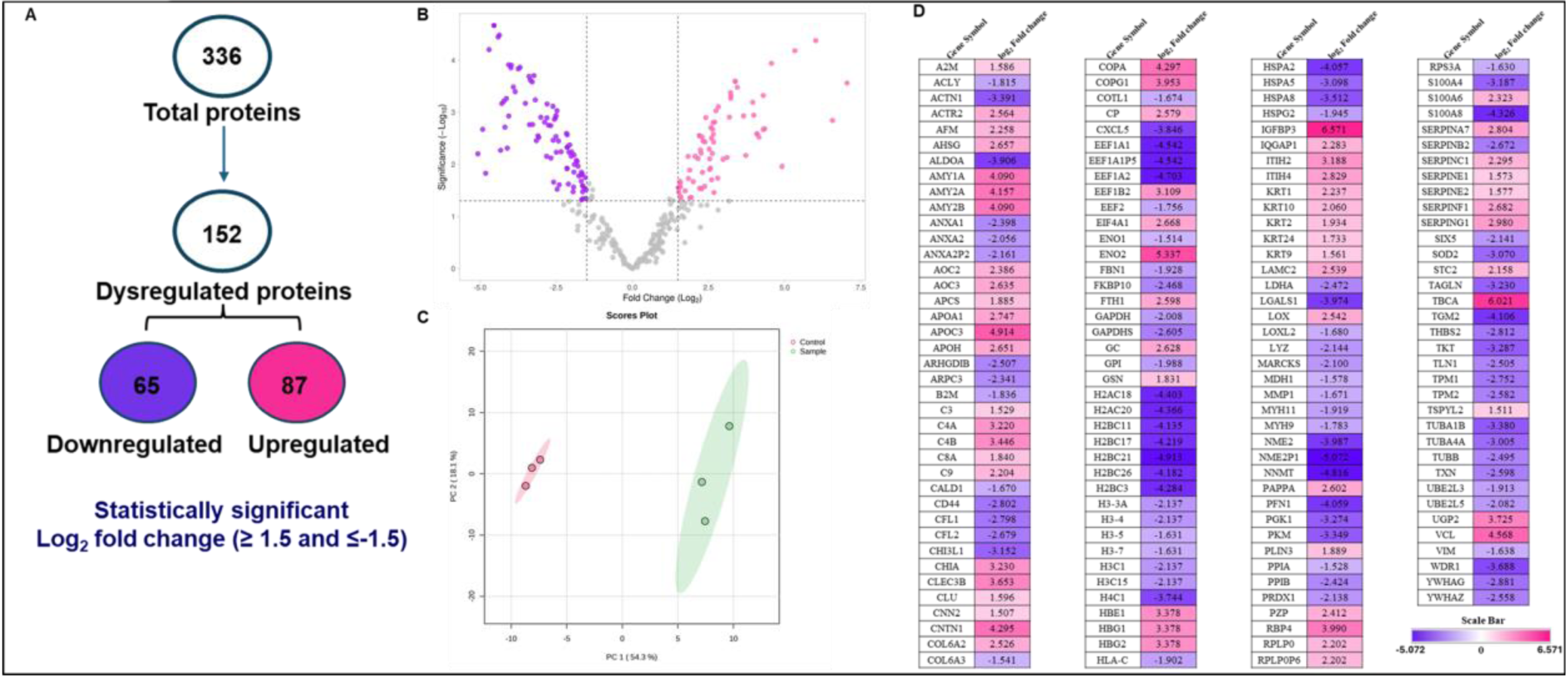
Label-free proteomics analysis of the CCM of WJ-MSCs and its direct co-culture with aPBMNCs using LC-MS/MS. (A) A flow chart depicts the total number of identified and dysregulated proteins. (B) A volcano plot highlights differentially expressed proteins. (C) Principal component analysis (PCA) demonstrates the good reproducibility of each biological replicate. (D) A heat map shows the expression levels of dysregulated proteins. Independent experiments were conducted with three different donors (biological replicates).

Further functional enrichment analysis of upregulated proteins revealed that these proteins were involved in plasma lipoprotein assembly (21.05%), carbohydrate metabolism (15.79%), and complement cascade regulation (26.32%) (Figure S13A-B). KEGG pathway analysis underscored the role of metabolic reprogramming in immune regulation, with significant enrichment of proteins associated with carbohydrate metabolism (33.33%) and cholesterol metabolism (16.67%) (Figure S13C). Immune system processes analysis of these upregulated proteins highlighted a significant enrichment of proteins involved in the complement system, including the lectin pathway (42.88%), alternative pathway (14.29%), and classical pathway (28.57%) (Figure S13D).

The functional enrichment analysis of downregulated proteins in the co-culture of WJ-MSCs-CCM revealed significant insights into immunomodulation and metabolic reprogramming. The Reactome pathway analysis indicated that a significant proportion of proteins were involved in the activation of RHO GTPases, which activate PKNs (83.52%), involved in cell growth and differentiation. Additionally, gluconeogenesis (3.3%) and gene and protein expression regulated by JAK-STAT signaling following interleukin-12 stimulation (3.3%) were also downregulated in the co-culture of WJ-MSCs (Figure S14A-B). In terms of pathway analysis, the KEGG pathways showed that a substantial proportion of the downregulated proteins were involved in systemic lupus erythematosus (SLE) (37.5%), gluconeogenesis (12.5%), and antigen processing and presentation (12.5%) (Figure S14C). Notably, 50.0% of the identified proteins were associated with the antimicrobial humoral immune response, specifically mediated by antimicrobial peptides (Figure S14D).

### WJ-MSCs^HYP+APO^-CCM versus CCM of coculture of WJ-MSCs^HYP+APO^ with aGVHD patients-derived aPBMNCs

Moreover, the comparative analysis of WJ-MSCs^HYP+APO^-CCM compared to CCM of its coculture with aGVHD patients-derived aPBMNCs revealed 369 total proteins. Among these, 83 proteins displayed statistically significant alterations with a Log_2_ fold change threshold of ≥ 1.5 and ≤ -1.5. Moreover, we observed that 39 dysregulated proteins were downregulated, while 44 were upregulated in the WJ-MSCs^HYP+APO^-CCM (Figure 9A-D).

**Figure 9:**
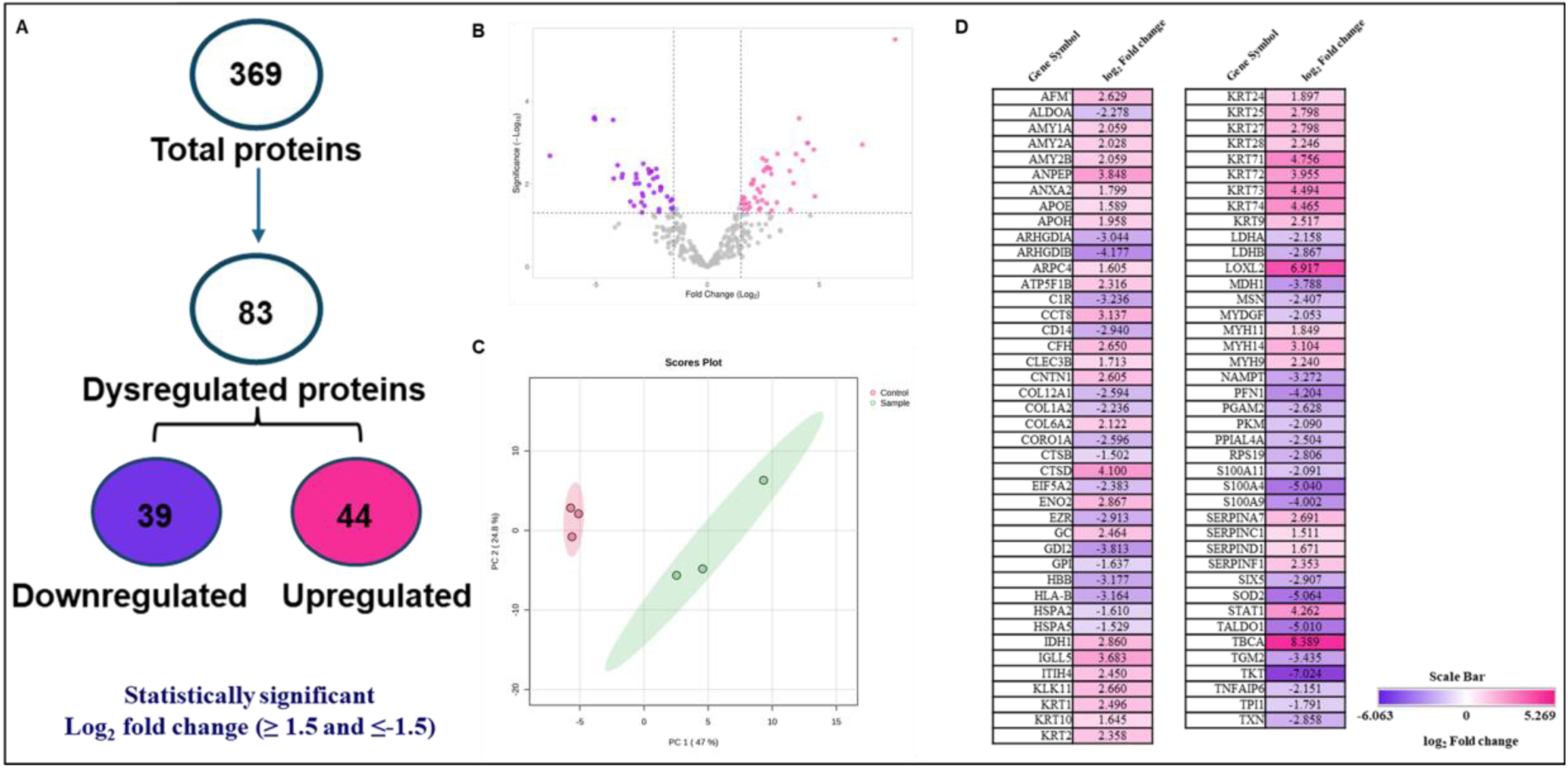
Label-free proteomics analysis of the CCM of WJ-MSCs^HYP+APO^ and its direct co-culture with aPBMNCs using LC-MS/MS. (A) A flow chart depicts the total number of identified and dysregulated proteins. (B) A volcano plot highlights differentially expressed proteins. (C) Principal component analysis (PCA) demonstrates the good reproducibility of each biological replicate. (D) A heat map shows the expression levels of dysregulated proteins. Independent experiments were conducted with three different donors (biological replicates).

Biological and molecular function analyses of the upregulated proteins revealed that WJ-MSCs^HYP+APO^-CCM mediate immune regulation and tissue repair through multiple mechanisms, including regulation of blood coagulation (73.08%), intermediate filament organization (11.54%), alpha-amylase activity (66.67%), and serine-type endopeptidase inhibitor (33.33%) (Figure S15A-B). Further cellular component analysis of these proteins revealed that a significant proportion of the proteins, accounting for 30.0%, were localized within the intermediate filament cytoskeleton and myosin II complex. Following this, 20.0% of the identified proteins were found to reside in the platelet-dense granule lumen, reflecting potential roles in hemostasis and immune regulation. Additionally, 10.0% of the proteins were linked to a cornified envelope (Figure S15C).

Further functional enrichment analysis of upregulated proteins revealed that these proteins were involved in RHO GTPase activating CIT (61.54%), and the digestion of dietary carbohydrate (23.08%) (Figure S16A-B). Wiki pathway analysis underscored the role of the complement system in immune regulation, with significant enrichment of proteins associated with the complement and coagulation cascade (100.0%) (Figure S16C). KEGG pathway analysis of these upregulated proteins highlighted a significant enrichment of proteins involved in starch and sucrose metabolism (50.0%) and *Staphylococcus aureus* infection (Figure S16D).

The functional enrichment analysis of downregulated proteins in the WJ-MSCs^HYP+APO^-CCM revealed significant insights into immunomodulation through metabolic reprogramming. The KEGG and Wiki pathways showed that a substantial proportion of the downregulated proteins were involved in gluconeogenesis (66.67%), aerobic glycolysis (66.67%), and antigen processing and presentation (16.67%) (Figure S17A-C). The Reactome pathway analysis indicated that a major proportion of proteins were involved in the exocytosis of proteins (33.33%) (Figure S17D).

### CCM of coculture of WJ-MSCs with aGVHD patients-derived aPBMNCs versus CCM of coculture of WJ-MSCs^HYP+APO^ with aGVHD patients-derived aPBMNCs

To investigate the mechanistic basis underlying the enhanced immunomodulatory potential of WJ-MSCs^HYP+APO^ observed *in vitro*, we performed proteomic analysis of the CCM from cocultures of WJ-MSCs or WJ-MSCs ^HYP+APO^ with aPBMNCs derived from aGVHD patients. Among 433 identified proteins, 95 were significantly dysregulated (log₂ fold change ≥ 1.5 or ≤ –1.5), with 52 proteins upregulated and 43 downregulated in the CCM derived from WJ-MSCs compared to WJ-MSCs^HYP+APO^ cocultured CCM (Figure 10A). The volcano plot (Figure 10B) and PCA plot (Figure 10C) illustrate the clear segregation and distribution of protein expression between the two groups, while the heat map (Figure 10D) highlights the clustering pattern of the dysregulated proteins.

**Figure 10:**
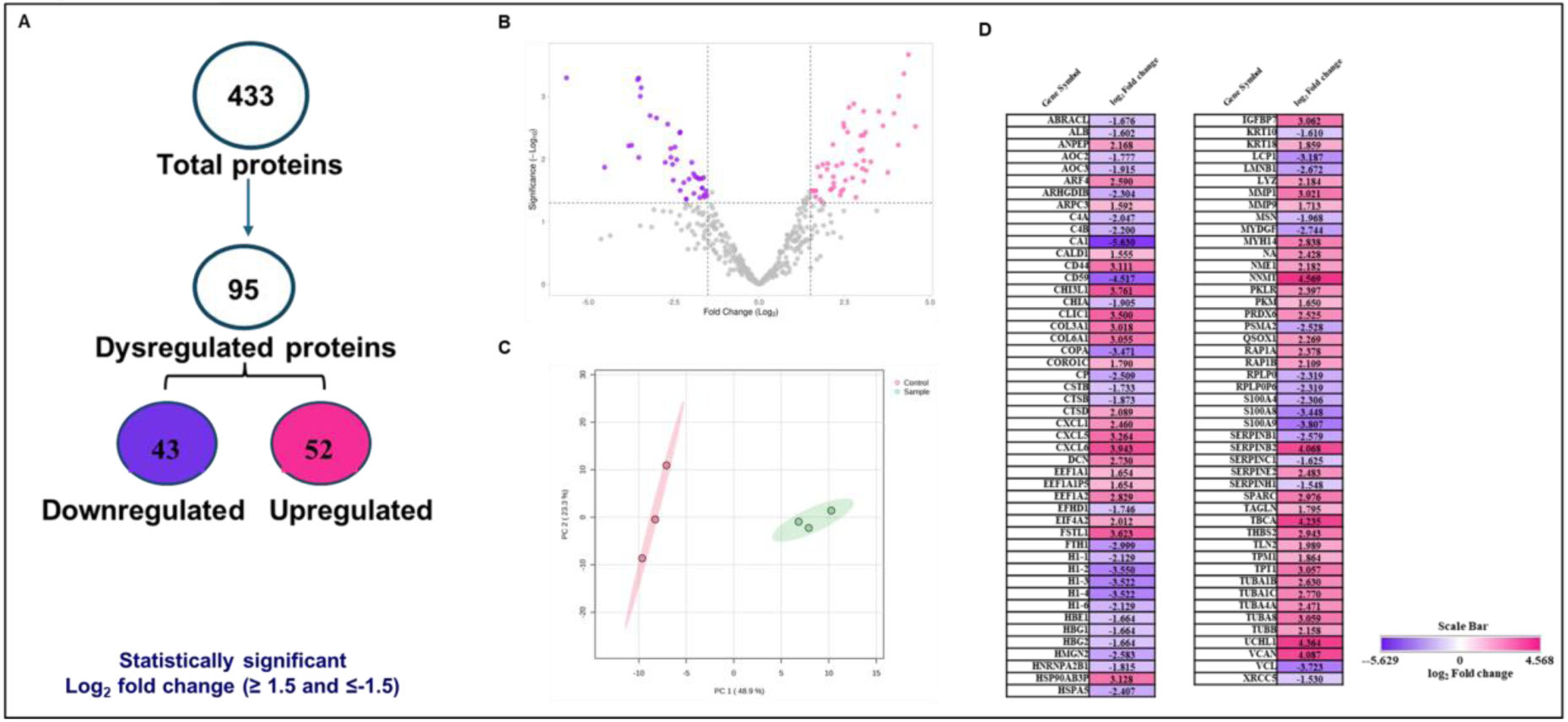
Label-free proteomics analysis of the CCM of coculture of WJ-MSCs with aGVHD patients-derived aPBMNCs and coculture of WJ-MSCs^HYP+APO^ with aPBMNCs derived from aGVHD patients using LC-MS/MS. (A) A flow chart depicts the total number of identified and dysregulated proteins. (B) A volcano plot highlights differentially expressed proteins. (C) Principal component analysis (PCA) demonstrates the good reproducibility of each biological replicate. (D) A heat map shows the expression levels of dysregulated proteins. Independent experiments were conducted with three different donors (biological replicates).

Gene Ontology (GO) enrichment of upregulated proteins in CCM of coculture of WJ-MSCs with aPBMNCs revealed enrichment in CXCR chemokine receptor binding (50.0%), collagen binding (16.67%), and translation elongation factor activity (33.33%) (Figure S18A). The biological processes included translation elongation (60.0%), chemokine activity (20.0%), and structural components of the cytoskeleton (20.0%) (Figure S18B). These proteins were predominantly localized to the tertiary granule lumen (50.0%), platelet alpha granule (25.0%), and ruffle membrane (25.0%) (Figure S18C). Pathway enrichment via Wiki and KEGG pathways linked these proteins to inflammatory and stress-associated processes such as pathogenic *E. coli* infection (25.0%), IL-17 signalling pathway (40.0%), tight junction remodelling (40.0%), Legionellosis (20.0%), hepatitis C and hepatocellular carcinoma (12.5%), burn wound healing (12.5%), and SARS-CoV-2 innate immunity evasion (12.5%). Reactome pathway analysis indicated involvement in post-chaperonin tubulin folding (79.49%) and ECM degradation (12.82%) (Figure S19A-D).

In contrast, downregulated proteins in WJ-MSCs coculture CCM, and thus upregulated in WJ-MSCs^HYP+APO^ coculture CCM, were enriched in functions highly relevant to immunomodulation and aGVHD mitigation. This included hemoglobin alpha binding (46.15%), nucleosomal DNA binding (23.08%), RAGE receptor binding (15.38%), complement binding (7.69%), and copper ion binding (7.69%) (Figure S20A). Biological processes represented among these proteins included antioxidant activity (40.0%), leukocyte aggregation (32.0%), negative regulation of DNA recombination (24.0%), and ribosome assembly (4.0%) (Figure S20B). These proteins localized predominantly to immunologically relevant compartments such as ficolin-1-rich granule lumen (25.0%), euchromatin (25.0%), blood microparticles (25.0%), and hemoglobin complexes (25.0%) (Figure S20C).

KEGG pathway revealed a striking downregulation (100%) of proteins involved in the complement and coagulation cascades, and reactome analysis showed involvement in the formation of senescence-associated heterochromatin foci (SAHF) (36.36%), complement activation (18.18%), post-translational phosphorylation (18.18%), and JAK-STAT signaling following IL-12 stimulation (27.27%) (Figure S21A-C).

Collectively, these proteomic findings corroborate the superior immunoregulatory profile of WJ-MSCs^HYP+APO^ observed *in vitro*. The downregulation of pro-inflammatory pathways and upregulation of antioxidant, complement-regulatory, and immune-calming pathways suggest that WJ-MSCs^HYP+APO^ exert a more potent modulatory effect on immune cells from aGVHD patients, offering a promising advantage over native WJ-MSCs in the context of aGVHD therapy.

## DISCUSSION

Our findings align with the existing literature supporting the use of MSCs as a therapeutic option in immunological disorders, notably aGVHD (25). Our results revealed that WJ-MSCs-CCM exhibit superior immunosuppressive capabilities compared to BM-MSCs-CCM, irrespective of naïve or preconditioned.

The observed inhibition of CD3^+^ T-cell proliferation by MSCs-CCM parallels existing literature, which demonstrated that MSCs can significantly suppress T-cell activation and proliferation in various immunological contexts through the paracrine mechanism (26,27). Our data indicating a greater inhibition of T-cell proliferation with WJ-MSCs compared to BM-MSCs reinforces earlier studies suggesting that WJ-MSCs possess unique properties that enhance their immunomodulatory effects, potentially due to their lower expression of MHC molecules and co-stimulatory signals, facilitating a more robust immunosuppressive response (28).

Hypoxia preconditioning has been shown to enhance immunosuppressive properties of MSCs, a finding consistent with recent research indicating that hypoxic conditions upregulate immunomodulatory factor expression in MSCs, thereby improving their therapeutic efficacy (29–31). The results here robustly demonstrate that both WJ-MSCs and BM-MSCs benefit from hypoxic preconditioning, suggesting a strategy to optimize MSC therapy through environmental manipulation before administration.

In terms of T-cell subset dynamics, our data showed a skewing of T-cell responses, with a reduction in CD8^+^ T-cell and an increase in the CD4^+^/CD8^+^ ratio following MSCs treatment, a significant observation considering that CD8^+^ T-cell is the primary effector cells in GVHD pathogenesis. This is in line with previous findings that reported MSCs can modulate the balance of T-helper cell subsets by promoting Tregs generation while inhibiting effector T-cell populations (32,33). Specifically, the ability of WJ-MSCs to enhance Tregs induction while simultaneously suppressing CD8^+^ T-cell proliferation suggests a dual mechanism through which they exert their immunomodulatory effects, promoting an anti-inflammatory state that can be crucial in aGVHD management. Furthermore, the significant increase in Tregs generation has been linked to better clinical outcomes in transplantation, indicating that targeting Tregs through MSCs-CCM could be an effective strategy to mitigate adverse immune responses.

Our results indicate that MSCs-CCM not only reduced the M1 macrophage phenotype but also promoted an M2 anti-inflammatory phenotype, as evidenced by increased Arginase-1 expression. This is consistent with prior research detailing how MSCs can polarize macrophages towards an M2 phenotype, which has been associated with enhanced tissue repair and resolution of inflammation (34). Our finding that WJ-MSCs-CCM outperforms BM-MSCs-CCM in promoting the M2 phenotype, irrespective of naïve or preconditioned state suggests an intrinsic property of WJ-MSCs that could leverage their clinical application in inflammatory conditions.

Additionally, mitochondrial function assays provided insights into the metabolic alterations that accompany MSCs-CCM treatment. The restoration of mitochondrial polarization indicates a positive metabolic shift towards oxidative phosphorylation, aligning with previous findings (35,36) that highlights the role of mitochondrial metabolism in maintaining T-cell function during immunosuppressive treatment strategies. The enhancement of T-cell mitochondrial health seen here underlies the immunomodulatory effects observed, providing a mechanistic link between MSCs-CCM treatment and improved T-cell functionality during aGVHD.

The *in vivo* results of our study demonstrate compelling evidence supporting the therapeutic potential of WJ-MSCs^HYP+APO^-CCM in the treatment of aGVHD. A major reduction in the incidence of diarrhea, skin ruffling, hunching, and improvement in the opening of the eyelid in the WJ-MSCs^HYP+APO^-CCM-treated groups, contrasting sharply with the aGVHD control group, provides critical insight into the protective effects of hypoxia-apoptosis conditioned MSCs on intestinal integrity. It has been well-documented that gastrointestinal symptoms and skin are prevalent in aGVHD and correlate with increased morbidity and mortality, reflecting the severity of the immune response against host tissues (37). Our findings indicated that WJ-MSCs^HYP+APO^-CCM effectively mitigates aGVHD symptoms, confirming earlier investigations, which demonstrated that MSC therapies can attenuate skin GVHD (38) and gastrointestinal inflammation (39) and improve overall clinical performance in aGVHD models. To the best of our knowledge, this is the first study, where we explored and compared the immunomodulatory effects of naïve and preconditioned MSCs-derived CCM in a preclinical aGVHD model.

Our *in vitro* and *in vivo* studies elucidate the immune restoration capabilities of WJ-MSCs (naïve, preconditioned) and their CCM in the context of aGVHD. The label-free proteomics analysis provided profound insights into the underlying mechanisms by which WJ-MSCs^HYP+APO^-CCM mediates immunomodulation compared to standard WJ-MSCs-CCM. The comparative evaluation sheds light on the complex interplay of immune and metabolic pathways influenced by MSCs-derived secretome.

Analysis revealed WJ-MSCs-CCM linked to crucial immunological functions, including a notable emphasis on processes such as immune regulation through protein processing and cellular responses to amino acid stimuli. This finding resonates with existing literature that underscores the impact of ECM components in modulating local immune responses, emphasizing the role of ECM-derived factors in tissue homeostasis and inflammation (40). The predominance of growth factor binding highlights the central role of signaling molecules in mediating functions of MSCs, as seen in studies demonstrating that MSCs promote tissue repair through growth factor release that enhances cell proliferation and survival (41,42).

The biological functions of the downregulated proteins provide additional layers of insight into the mechanism by which WJ-MSCs exert their effects. The significant enrichment of downregulated proteins linked to nucleosome assembly and phospholipase A2 inhibitor activity is particularly compelling, as it correlates with findings that highlight the influence of nucleosome dynamics on chromatin remodeling in inflammatory diseases (43). Enzymes involved in the production of phospholipid mediators have been shown to modulate inflammation, impacting leukocyte recruitment and activation, which is a pivotal aspect of aGVHD pathophysiology.

The proteomics analysis of WJ-MSCs^HYP+APO^-CCM alone and their co-culture with aGVHD patients derived aPBMNCs immunomodulatory properties of MSCs influenced by their cellular environment, leading to a differential expression of factors conducive to immune regulation. Pathway analyses underscored the importance of metabolic shifts, as evidenced by significant enrichment in proteins associated with plasma lipoprotein assembly and cholesterol metabolism, highlighting their role in modulating immune responses and supporting earlier conclusions that metabolic alterations directly influence T-cell activity (35,36). MSCs can positively regulate local immune responses through complement pathway engagement, consistent with the previous findings (44,45). This is especially relevant considering the role of complement proteins in enhancing phagocytic activity and shaping adaptive immune responses, demonstrating MSCs’ capabilities as potent modulators of the immune landscape.

Collectively, our findings draw compelling connections between WJ-MSCs and broader immunological and metabolic paradigms. The potential of WJ-MSCs^HYP+APO^-CCM to downregulate pathways associated with gluconeogenesis and enhance those linked to ECM functions emphasizes their multifaceted role in promoting immune tolerance and limiting tissue damage during aGVHD. Taken together, these results advance our understanding of MSCs-derived CCM as a valuable therapeutic approach for aGVHD and set the stage for further investigation into specific molecular mechanisms, potentially facilitating the development of targeted MSCs-based therapies.

## CONCLUSION

In conclusion, WJ-MSCs^HYP+APO^-CCM demonstrate significant immunomodulatory capabilities, highlighting their role in promoting immune restoration while providing insights into metabolic shifts critical for successful outcomes in aGVHD management. These findings contribute substantially to the development of effective MSCs-based therapeutic interventions, advancing the field of immunomodulation.

## MATERIAL AND METHODS

### Sex as a biological variable

Our study involved both male and female human subjects. Efforts were made to recruit participants of both sexes, and data were analyzed without prior assumptions regarding sex-based differences. However, the sample size was not powered to detect sex-specific effects. Therefore, while the findings are expected to be broadly relevant across sexes, potential sex-dimorphic responses cannot be ruled out. Additionally, 10– 12 weeks old male C57BL/6 (H-2K^b^) donor and female BALB/c (H-2K^d^) recipient mice, weighing 24±2 grams, were selected for our study.

### Cell lines

The Human GFP-tagged immortalized human fibroblast (IHF), MOLM-13, and Kasumi-1 cell lines were kindly gifted by the All India Institute of Medical Sciences (AIIMS), New Delhi, India. IHF cells were maintained in 1X HG-DMEM (Thermo Fisher Scientific, USA) supplemented with 10% FBS and 1% Antibiotic-Antimycotic solution (Thermo Fisher Scientific, USA). MOLM-1 and Kasumi-1 cells were maintained in RPMI-1640 medium supplemented with 10% FBS (Thermo Fisher Scientific, USA) and 1% Antibiotic-Antimycotic solution (Thermo Fisher Scientific, USA).

### Isolation and preconditioning of MSCs from human bone marrow (BM) and Wharton’s Jelly (WJ)

The study involving human subjects and the use of human MSCs was approved by the Institutional Human Ethics Committee (Ref. No. IECPG-542/23.09.2020) and the Institutional Committee for Stem Cell Research at the same institution [Ref. No.: IC-SCR/110/20(R)] respectively at the AIIMS, New Delhi, India. Informed written consent was obtained from all participants, and all procedures were conducted following the guidelines and regulations approved by the ethics committee.

Human bone marrow (BM) aspirates were obtained from ten healthy donors who were recipients of patients undergoing allogeneic stem cell transplantation at the Department of Medical Oncology, Dr. B. R. Ambedkar Institute Rotary Cancer Hospital, AIIMS, New Delhi. Additionally, human umbilical cord (UC) tissue was collected from ten donors in sterile transport media containing 1000 IU/ml heparin and 200 μg/ml gentamicin from the Department of Obstetrics and Gynaecology at AIIMS, New Delhi. MSCs from both BM and Wharton’s Jelly (WJ) were isolated following our established protocol (7). Passages 3-5 were used for subsequent *in vitro* experiments, and these were pooled together at their respective passages.

MSCs were preconditioned with 1% O2 in 1X LG-DMEM complete media for 24 hours in a tri-gas incubator (Thermo Fisher Scientific, USA), termed hypoxia-preconditioned MSCs (MSCs^HYP^). Apoptotic MSCs (MSCs^APO^) were generated using 0.5µM staurosporine (STS) (Sigma, USA) in 1X LG-DMEM complete media in a humidified chamber at 37°C with 5% CO_2_ for 24 hours. Hypoxia-preconditioned apoptotic MSCs (MSCs^HYP+APO^) were generated by exposing them to 1% O_2_ and 1µM STS for 24 hours in a tri-gas incubator (7).

### Collection of MSCs-CCM

Before collecting the CCM from each group (MSCs, MSCs^HYP^, MSCs^APO^, MSCs^HYP+APO^), the fetal bovine serum (FBS)-containing culture medium was replaced with Stem Pro™ MSC serum-free media (SFM) (Thermo Fisher Scientific, USA) to eliminate xenogenic components. After 48 hours, the CCM was collected and centrifuged at 3,500 rpm for 10 minutes. The supernatant was then filtered through a 0.2 μm syringe filter (Sigma-Aldrich, Saint Louis, MO, USA) to remove cell debris and stored at -80 °C for further experiments.

### *In vitro* T-cell proliferation assay

Peripheral blood (PB) was collected from grade II-IV aGvHD patients (n=25) using sterile sodium heparin-coated vacutainers (BD Biosciences, US). The isolation of peripheral blood mononuclear cells (PBMNCs) was performed through Ficoll density gradient centrifugation (8).

Subsequently, CD3^+^ T-cell were extracted from the PBMNCs through negative selection using a Pan T cell isolation kit (Miltenyi Biotec, USA), following the manufacturer’s protocol. The isolated CD3^+^ T-cell were stained with 1 µM cell Trace^TM^ carboxyfluorescein succinimidyl ester (CFSE) dye (BD Biosciences, USA) and activated with PHA (1µg/ml) (Sigma, USA) and IL-2 (50IU/ml) (Thermo Fisher Scientific, USA) using our standardized protocol (7).

Moreover, CFSE-labelled activated T-cell were treated with MSCs-CCM (MSCs, MSCs^HYP^, MSCs^APO^, MSCs^HYP+APO^) and 1X RPMI-1640 complete medium (Thermo Fisher Scientific, USA) in a 1:1 ratio for 3 days.

After the treatment, T-cell proliferation was evaluated using a DxFlex flow cytometer (Beckman Coulter, USA), and the data analysis was performed using Kaluza software version 2.1 (Beckman Coulter, USA). The normalization of T-cell proliferation was performed with CFSE-labeled activated T-cell only (7).

### Induction of regulatory T-cell (Tregs)

Activated T-cell were treated with MSCs-CCM (MSCs, MSCs^HYP^, MSCs^APO^, MSCs^HYP+APO^) and 1X RPMI-1640 complete medium (Thermo Fisher Scientific, USA) in a 1:1 ratio for 5 days. Following this period, the cells were collected, washed with 1X PBS, and stained with fluorochrome-conjugated anti-human monoclonal antibodies targeting CD3, CD4, CD8, CD25, and CD127 (Beckman Coulter, USA). A minimum of 50,000 events was acquired using a DxFlex flow cytometer (Beckman Coulter, USA), and the data were analyzed with Kaluza software version 2.1 (Beckman Coulter). Activated T-cell cultured without MSCs served as a control to establish baseline expression levels of CD3^+^ CD4^+^ CD25^+^ FOXP3^+^ Tregs (7,9).

### Enumeration of effector memory helper T cell subtypes (Th1, Th2, and Th17)

The proportions of Th1, Th2, and Th17 cells were quantified following 5-day exposure of activated T-cell to MSCs-CCM (MSCs, MSCs^HYP^, MSCs^APO^, MSCs^HYP+APO^), using surface staining with fluorochrome-conjugated anti-human monoclonal antibodies against CXCR3, CXCR5, CCR10, CCR4, CCR7, and CCR6 at 37°C for 30 minutes. This was followed by additional staining with anti-human fluorochrome-conjugated monoclonal antibodies specific for CD3, CD4, CD8, and CD45RA. A minimum of 50,000 cells was acquired using a DxFlex flow cytometer (Beckman Coulter, USA), and the data were analyzed using Kaluza software version 2.1 (Beckman Coulter, USA). Activated T-cell in the absence of MSCs-CCM served as a control to establish baseline expression levels of Th1, Th2, and Th17 (7).

### Enumeration of αβ and γδ CD4^+^ T-cell

The proportions of αβ and γδ CD4^+^ T-cell was quantified in the 3-day culture of activated T-cell with MSCs-CCM (MSCs, MSCs^HYP^, MSCs^APO^, MSCs^HYP+APO^) by staining with fluorochrome-conjugated anti-human monoclonal antibodies specific for CD3, CD4, CD8, αβ, and γδ monoclonal antibodies for 30 minutes in the dark. A minimum of 50,000 cells was acquired using a DxFlex flow cytometer (Beckman Coulter, USA), and the data were analyzed using Kaluza software version 2.1 (Beckman Coulter, USA). Activated T-cell cultured without MSCs served as a control to establish baseline expression levels of αβ and γδ T-cell (10).

### Macrophage polarization

CD14^+^ monocytes were isolated from PBMNCs using a pan-monocyte isolation kit (Miltenyi Biotec, USA), following the manufacturer’s instructions. The isolated monocytes were differentiated into M1 macrophages using our previously established protocol (7). M1 macrophages were treated with MSCs-CCM (MSCs, MSCs^HYP^, MSCs^APO^, MSCs^HYP+APO^) and 1X RPMI-1640 complete medium (Thermo Fisher Scientific, USA) in a 1:1 ratio for 3 days. Cells were fixed and permeabilized using an IntraPrep permeabilization kit (Beckmann Coulter, USA), followed by intracellular staining of cells with fluorochrome-conjugated anti-human iNOS, Arginase-I (Thermo Fisher Scientific, USA) for 30 minutes. The cells were acquired using a DxFlex flow cytometer (Beckman Coulter, USA) to enumerate the M1 and M2 macrophages (7).

### Mitochondrial Reactive Oxygen Species (ROS) assay

The effect of MSCs-CCM on mitochondrial ROS of T-cell was assessed after the 3-days exposure of MSCs-CCM (MSCs, MSCs^HYP^, MSCs^APO^, MSCs^HYP+APO^) to activated T-cell. Following the treatment period, the cell suspension was stained with MitoSOX Red (Thermo Fisher Scientific, USA) at a concentration of 5μM at 37֯C for 20 minutes. Subsequently, the cells were washed with 1XPBS (Thermo Fisher Scientific, USA) and stained with fluorochrome-conjugated anti-human CD45 monoclonal antibody. The percentage of CD45^+^ MitoSox Red^+^ cells was quantified to enumerate the mitochondrial ROS level in aPBMNCs using a DxFlex flow cytometer (Beckmann Coulter, USA), and the data were analyzed using Kaluza Software Version 2.1. Activated T-cell cultured without CCM served as a control to establish baseline expression of mitochondrial ROS (11).

### Measurement of mitochondrial membrane potential

The mitochondrial health of T-cell was assessed after 3-day exposure of MSCs-CCM to activated T-cell by staining the cell suspension with JC-1 dye (Thermo Fisher Scientific, USA) at a concentration of 2μM at 37֯C for 20 minutes in the dark. Subsequently, the cells were washed with 1XPBS (Thermo Fisher Scientific, USA) and stained with fluorochrome-conjugated anti-human CD45 monoclonal antibody. The percentage of JC-1^+^ cells gated on CD45^+^ was enumerated to assess their mitochondrial health status (12).

### Sea-horse assay

Oxygen consumption rate (OCR) and extracellular acidification rate (ECAR) were assessed using the Seahorse XFe24 Extracellular Flux Analyzer (Agilent Technologies, California, USA), which serves as indicators for oxidative phosphorylation (OXPHOS) and glycolysis, respectively. T-cell were seeded at a density of 5*10^4^ cells per well, followed by the treatment of MSCs-CCM (MSCs, MSCs^HYP^, MSCs^APO^, MSCs^HYP+APO^) in a 1:1 ratio with 1X RPMI-1640 complete media (Thermo Fisher Scientific, USA) for 72 hours. The OCR measurements were taken sequentially, starting from the basal level, followed by the addition of 1.0 mM oligomycin, 0.75 mM FCCP (fluorocarbonyl cyanide phenylhydrazone), and a combination of 0.5 mM rotenone and antimycin to evaluate changes in mitochondrial respiratory parameters. Data analysis was performed using Wave 2.6.1 and GraphPad Prism software (13).

### Tube formation assay

The angiogenic potential of MSCs-CCM (WJ-MSCs-CCM and WJ-MSCs^HYP+APO^-CCM) was assessed using a tube formation assay on human umbilical vein endothelial cells (HUVECs) subjected to oxidative stress. Briefly, HUVECs were pre-exposed to 100 µM hydrogen peroxide (H₂O₂) for 2 hours to induce injury, followed by gentle washing with PBS. A 96-well plate was coated with 50 µl of Geltrex (Thermo Fisher Scientific, USA) per well and incubated at 37 °C for 1 hour to allow polymerization. The injured HUVECs were then seeded onto the Geltrex-coated wells, and either WJ-MSCs-CCM or WJ-MSCs^HYP+APO^-CCM was added simultaneously. After 24 hours, tube formation was visualized under an inverted microscope. The angiogenic parameters, including the number of nodes, meshes, and total tube length, were quantified using the Angiogenesis Analyzer plugin in ImageJ software (14).

### Cytotoxicity assay

The proliferation of Kasumi-1 and MOLM-13 cells after the treatment of MSCs-CCM (BM-MSCs, BM-MSCs^HYP+APO^, WJ-MSCs, WJ-MSCs^HYP+APO^) was assessed using the 3-(4,5-dimethylthiazol-2-yl)-5-(3-carboxymethoxyphenyl)-2-(4-sulfophenyl)-2H-tetrazolium (MTS) (Abcam, USA) following the manufacturer’s protocol. Briefly, 10μl of MTS (Abcam, USA) solution was added directly to each well of 96-well plates after 48 hours of treatment of MSCs-CCM. The plates were then incubated for 3 hours at 37֯C. Following incubation, absorbance was measured at 490 nm using an ELISA reader (Bio-Rad, USA), with the blank consisting of 1X RPMI-1640 medium culture medium alone (7).

The percentage of cell viability was calculated using the following formula:

Cell viability (%) = (Absorbance_treated_-Absorbance_blank_/Absorbance_control_-Absorbance_blank_) *100

### Experimental Animals

The study involved the use of mice and was approved by the Institutional Animal Ethics Committee, All India Institute of Medical Sciences, New Delhi (Ref. No. 358/IAEC-1/2022). 10–12 weeks old male C57BL/6 (H-2K^b^) donor and female BALB/c (H-2K^d^) recipient mice, weighing 24±2 grams were selected for our study. Mice were kept in the animal facility, where they had access to standard chow and water freely. They were maintained under a 12-hour light/dark cycle, with the environment set to a temperature range of 21-23°C and a relative humidity of 50%.

### Establishment of chemotherapy-based aGVHD murine model and MSCs and their CCM administration

The mismatched model of aGVHD was established using a chemotherapy approach. The chemotherapy model involved the intraperitoneal treatment of BALB/c recipient mice with busulfan (Sigma, USA) at a dosage of 20 mg/kg/day for four days (from D-7 to D-4), followed by cyclophosphamide (Sigma, USA) at 100 mg/kg/day for two days (from D-3 to D-2). A resting day was designated on D-1, during which no conditioning regimen doses were given. On day 0, the mice received 20*10^6^ splenic cells and 5*10^5^ bone marrow cells from C57BL/6 donor mice intravenously (15,16).

Upon the onset of aGVHD, the aGVHD mice were treated with intravenous injections of either 1*10^6^ MSCs (WJ-MSCs, WJ-MSCs^HYP+APO^) or 200 μl of their respective CCM (17). Two additional doses of MSCs or CCM were administered at the same dosage, with a two-day interval between each dose. Mice were monitored daily for clinical signs and symptoms indicative of aGVHD.

### Evaluation of donor cells engraftment

The engraftment of donor-positive cells (H-2K^b^) was evaluated using flow cytometry. Briefly, peripheral blood was collected from recipient BALB/c mice through the retro-orbital plexus. Following blood collection, red blood cells (RBCs) were lysed and washed thrice, and cells were stained with fluorochrome-conjugated anti-mouse monoclonal antibodies against H-2K^b^ MHC class I and H-2K^d^ MHC class I (Thermo Fisher Scientific, USA). A minimum of 10,000 cells was acquired using a DxFlex flow cytometer (Beckman Coulter, USA), and the data were analyzed using Kaluza software version 2.1 (Beckman Coulter, USA) (18).

### End-point analysis

The clinical aGVHD score, body weight, and overall survival were assessed in aGVHD murine models post MSCs and their CCM administration at regular intervals (D+5, D+12, D+19, D+26, D+40). The clinical scoring system was based on the following six criteria: fur texture, skin integrity, posture, activity, weight loss, and diarrhea to assess aGVHD severity (18,19).

### Histopathology

To assess the effect of MSCs and their CCM on target organ protection from aGVHD, target organs, namely the liver, lung, intestine, and skin, were harvested from the aGVHD control and treated groups and fixed in 10% neutral buffer formalin. The slides were stained with haematoxylin and eosin (H&E) in the animal histopathology laboratory at AIIMS. The tissue pathology evaluation and scoring were done as per the method reported earlier (20–22). The same H&E slides were used to score the lymphocytic infiltration in the GVHD target organs.

### *In vivo* serum cytokine quantification

The blood was collected from the retro-orbital sinus of both aGVHD control and treated mice. The serum levels of cytokines (IL-10, IFN-γ, IL-17α) were assessed using the ELISA as per the manufacturer’s instructions (Thermo Fisher Scientific, USA) (23).

### Identification of secreted proteins using liquid chromatography-assisted mass spectrometry (LC-MS/MS)

Protein per sample (WJ-MSCs, WJ-MSCs^HYP+APO^, and their co-culture with aPBMNCs) was used for digestion and reduced with 5 mM tris(2-carboxyethyl) phosphine (TCEP) and further alkylated with 50 mM iodoacetamide and then digested with Trypsin (1:50, Trypsin/lysate ratio) for 16 h at 37 °C. Digests were cleaned using a C18 silica cartridge to remove the salt and dried using a speed vac. The dried pellet was resuspended in buffer A (2% acetonitrile, 0.1% formic acid).

Experiments were performed on an Easy-nlc-1000 system coupled with an Orbitrap Exploris mass spectrometer. 1µg of peptide sample was loaded on C18 column 15 cm, 3.0μm Acclaim PepMap (Thermo Fisher Scientific, USA) and separated with a 0– 40% gradient of buffer B (80% acetonitrile, 0.1% formic acid) at a flow rate of 500 nl/min) and injected for MS analysis. LC gradients were run for 110 minutes. MS1 spectra were acquired in the Orbitrap (Max IT = 60ms, AGQ target = 300%; RF Lens = 70%; R=60K, mass range: 375−1500; Profile data). Dynamic exclusion was employed for 30s, excluding all charge states for a given precursor. MS2 spectra were collected for the top 20 peptides. MS2 (Max IT= 60ms, R= 15K, AGC target 100%). All samples were processed, and the RAW files generated were analyzed with Proteome Discoverer (v2.5) against the UniProt Human database. For dual Sequest and Amanda search, the precursor and fragment mass tolerances were set at 10 ppm and 0.02 Da, respectively. The protease used to generate peptides, i.e., enzyme specificity, was set for trypsin/P (cleavage at the C terminus of “K/R: unless followed by “P”). Carbamidomethyl on cysteine as fixed modification and oxidation of methionine and N-terminal acetylation were considered as variable modifications for database search. Both peptide spectrum match and protein false discovery rate were set to 0.01 FDR (24).

### Statistical analysis

All statistical analyses were conducted using GraphPad Prism version 8.4.3. One-way and Tukey’s post hoc tests compared three or more groups. Survival data were analyzed using the Kaplan–Meier method and Mantel-Cox log-rank test. Data was shown as Mean±S.D. and a p-value of ≤ 0.05 was considered statistically significant.

### Study approval

The study involved human subjects, which was approved by the Institutional Human Ethics Committee at the All India Institute of Medical Sciences, New Delhi, India (Ref. No.: IECPG-542/23.09.2020). Additionally, it included the use of human MSCs, which the Institutional Committee approved for Stem Cell Research at the same institution [Ref. No.: IC-SCR/110/20(R)]. Informed written consent was obtained from all participants, and all procedures were conducted in accordance with the guidelines and regulations approved by the ethics committee.

### Data availability

The data that support the findings of this study are available from the corresponding author upon reasonable request.

## AUTHOR’S CONTRIBUTIONS

MM performed the experiments, acquired, analyzed, and interpreted the data, and wrote the manuscript. MM was involved in performing experiments and data interpretation, and analysis. SR, RG was involved in data interpretation and analysis. SB, VD, DP, PSM, RP, MA, AKG, RD, TS, and MM provided patient samples and their clinical details. BN, TDS, SK, and RAM contributed to data interpretation and analysis. LM, SB, and GH were involved in the interpretation of proteomics data. HP and SM contributed to data interpretation, analysis, provided resources, and supervised the study. RKS conceptualized the study, provided resources, designed and supervised the experiments, interpreted the data, and reviewed and edited the manuscript. All authors critically reviewed and approved the final version of the manuscript.

## ACKNOWLEDGEMENT

The authors express their gratitude to the All India Institute of Medical Sciences (AIIMS), New Delhi, India, for facilitating the execution of the study. Schematic representative figures illustrating the methodology were created using Biorender.com.

## SUPPLEMENTARY FIGURES

**Figure S1:**
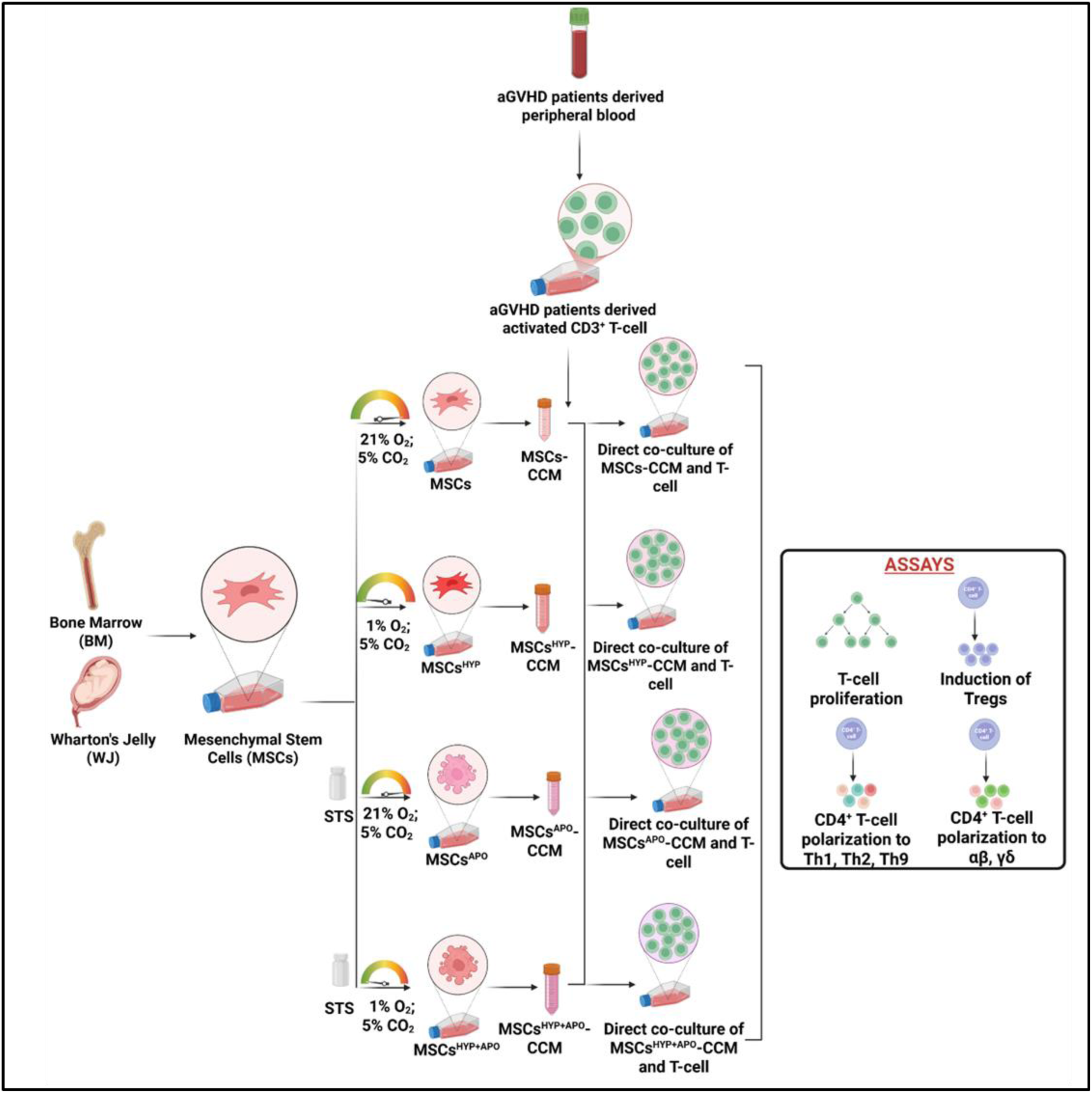
Schematic representation of the in vitro assays for the assessment of T-cell programming (created using Biorender.com).

**Figure S2:**
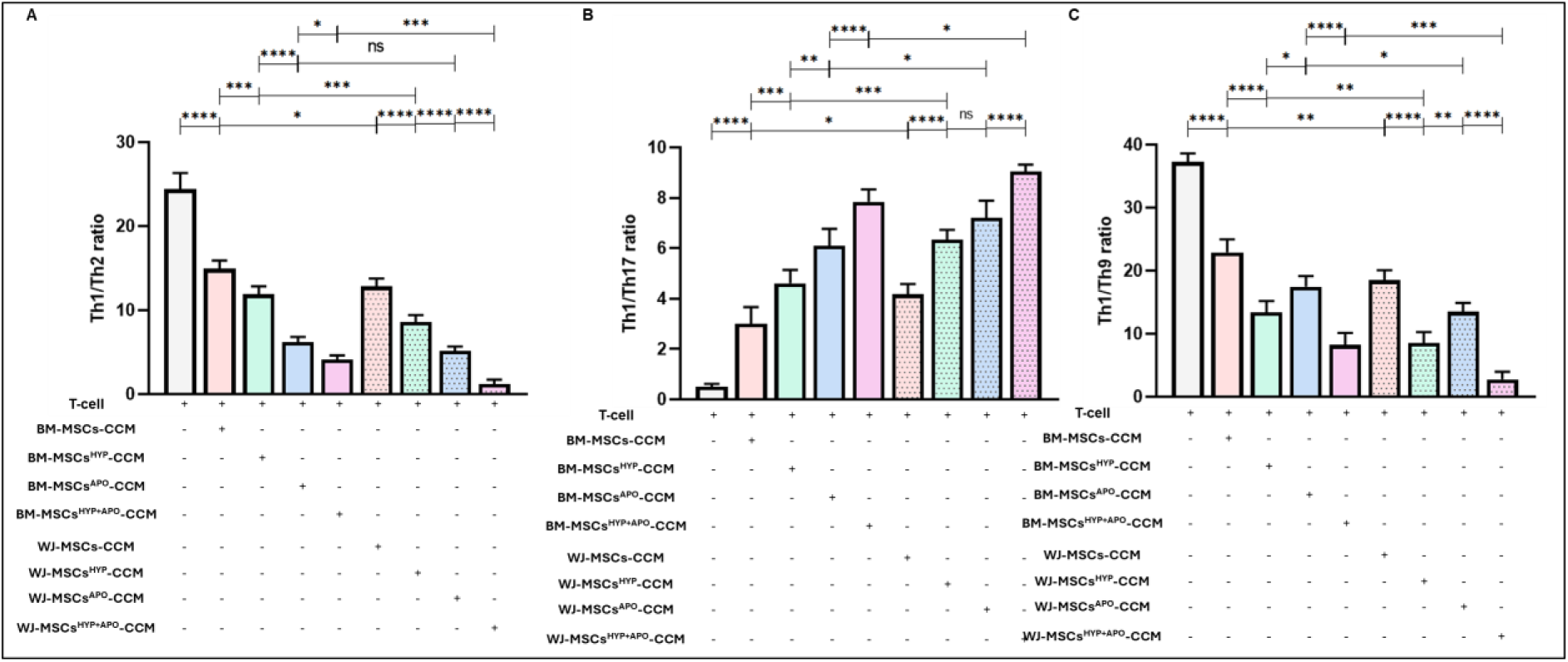
Effect of naïve, hypoxia, apoptotic, and hypoxia preconditioned apoptotic MSCs-CCM (MSCs, MSCs^HYP^, MSCs^APO^, MSCs^HYP+APO^) on the immunomodulatory reprogramming of T-cell derived from aGVHD patients. The bar graph represents (A) the ratio of Th1/Th2 (n=25). (B) the ratio of Th1/Th17 (n=25). (C) the ratio of Th1/Th9 (n=25) in the culture of MSCs-CCM and T-cell. Data shown represent the Mean±S.D of 25 independent experiments performed with T-cell derived from 25 different donors (biological replicates), with each experiment conducted in triplicate (technical replicates). Statistical analysis: Tukey’s multiple comparisons test; *≤0.05; **≤0.01; ***≤0.001; ****≤0.0001. Abbreviations: BM: Bone marrow; WJ: Wharton’s Jelly; MSCs: Mesenchymal Stem Cells; HYP: Hypoxia; APO: Apoptosis; CCM: Culture Conditioned Media

**Figure S3:**
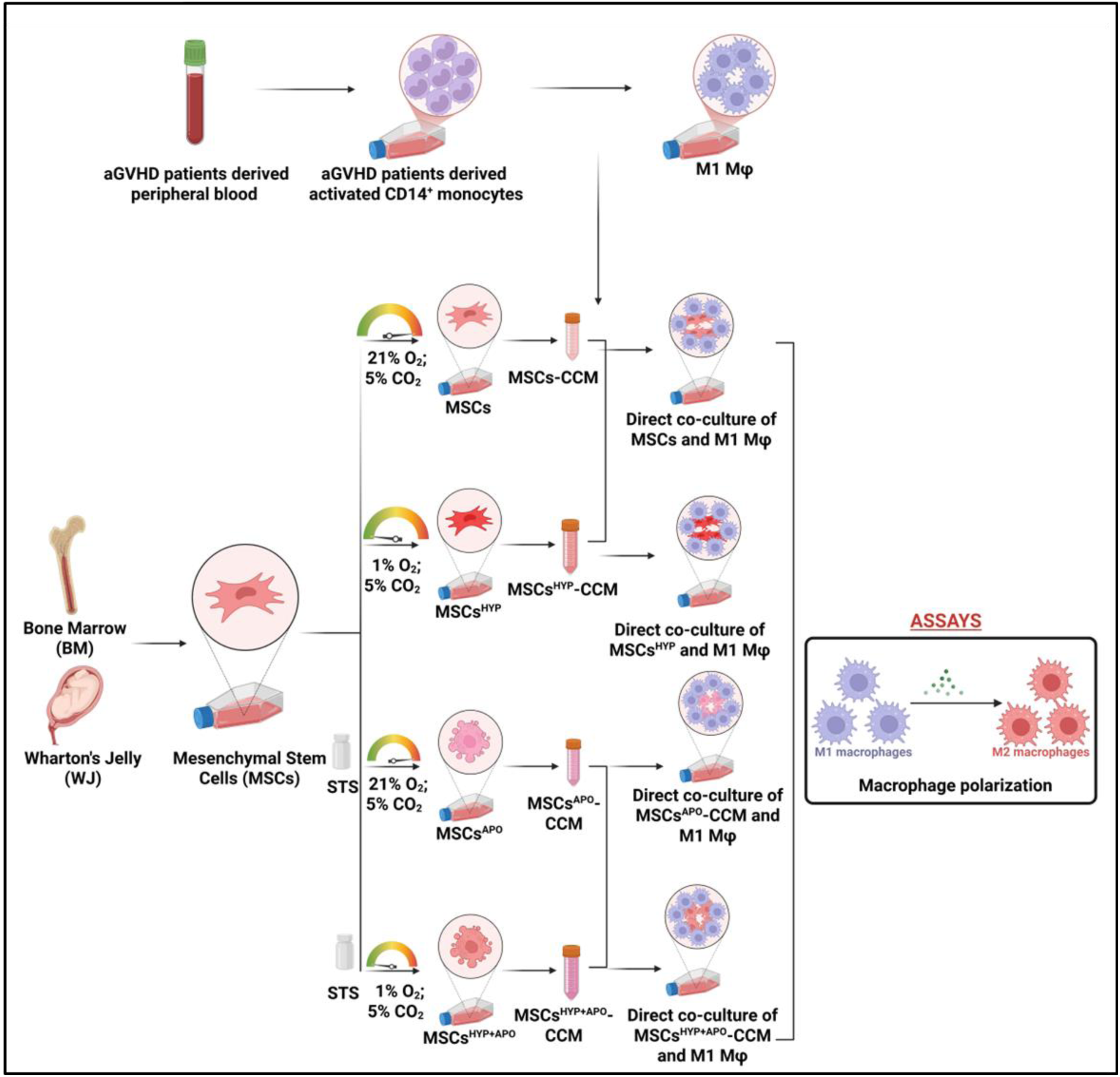
Diagrammatic representation of the assessment of the macrophage polarization (created using Biorender.com).

**Figure S4:**
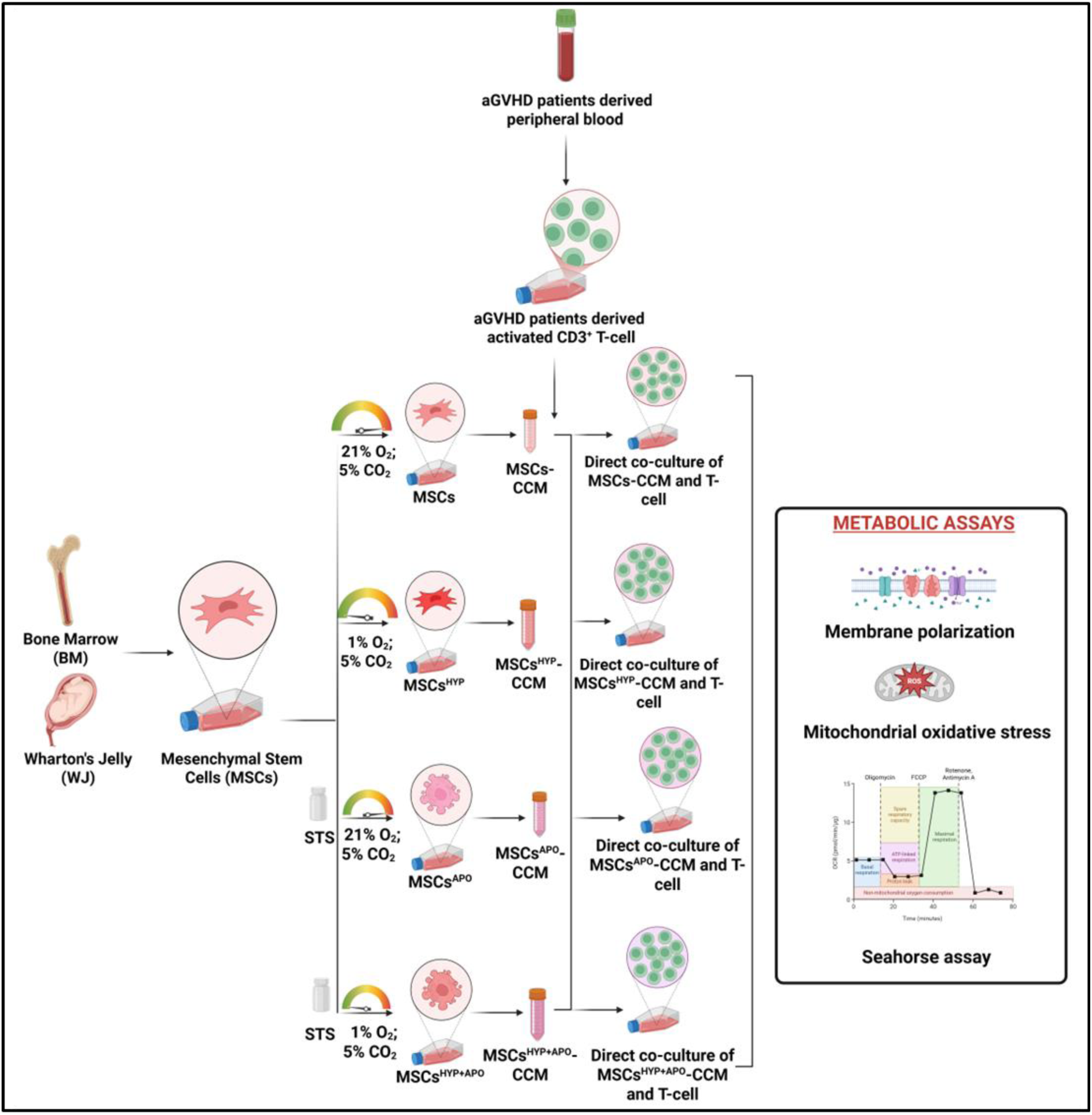
Diagrammatic representation for the assessment of modulation of T-cell metabolism and oxidative stress (created using Biorender.com).

**Figure S5:**
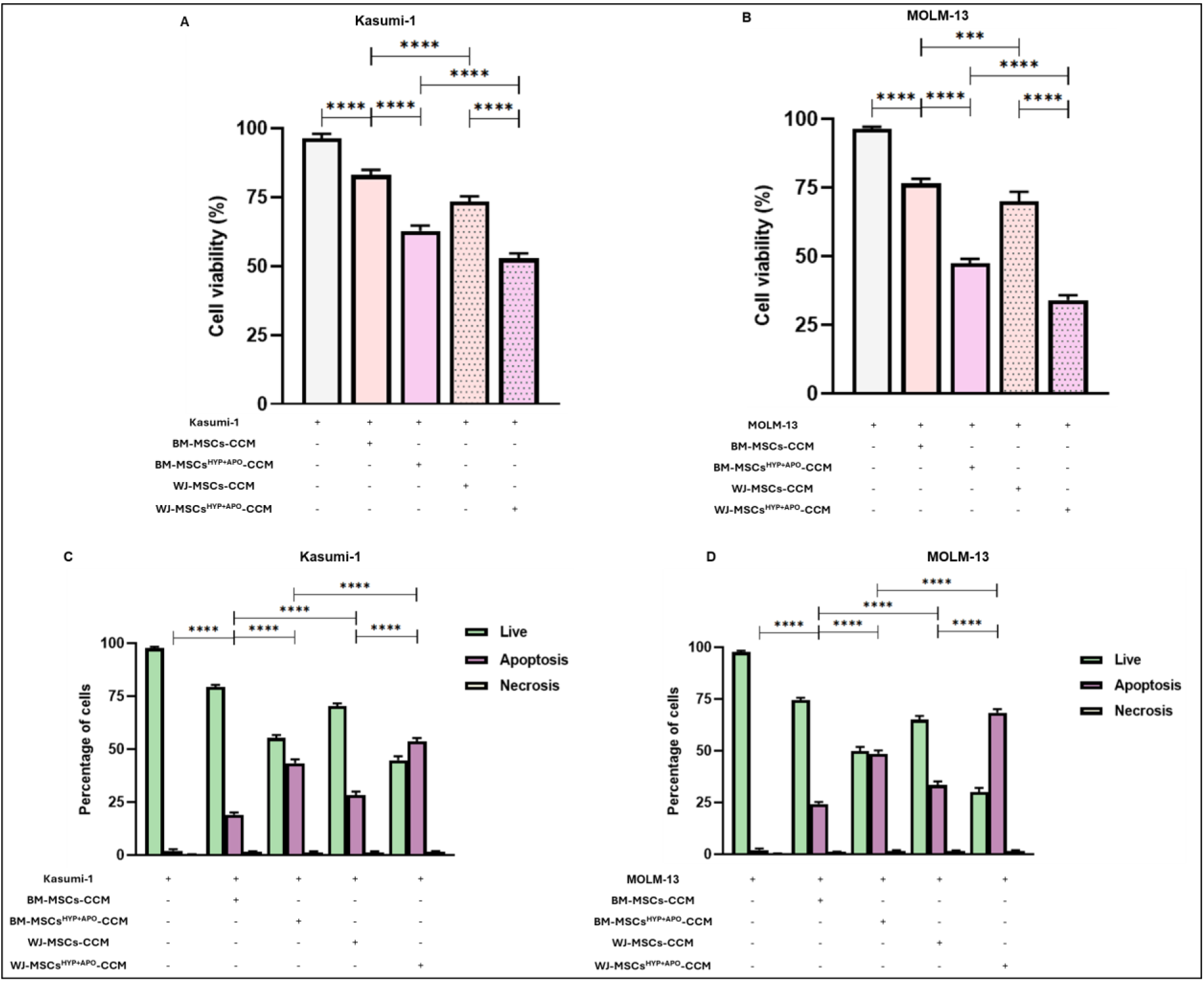
Effect of naïve and hypoxia preconditioned apoptotic MSCs-CCM (MSCs, MSCs^HYP+APO^) on the leukemic cell lines. The bar graph represents (A) cell viability (%) of Kasumi-1. (B) cell viability (%) of MOLM-13. (C) percentage of cells of Kasumi-1. (D) percentage of cells of MOLM-13. Data shown represent the Mean±S.D of 5 independent experiments, with each experiment conducted in triplicate (technical replicates). Statistical analysis: Tukey’s multiple comparisons test; *≤0.05; **≤0.01; ***≤0.001; ****≤0.0001. Abbreviations: BM: Bone marrow; WJ: Wharton’s Jelly; MSCs: Mesenchymal Stem Cells; HYP: Hypoxia; APO: Apoptosis; CCM: Culture Conditioned Media

**Figure S6:**
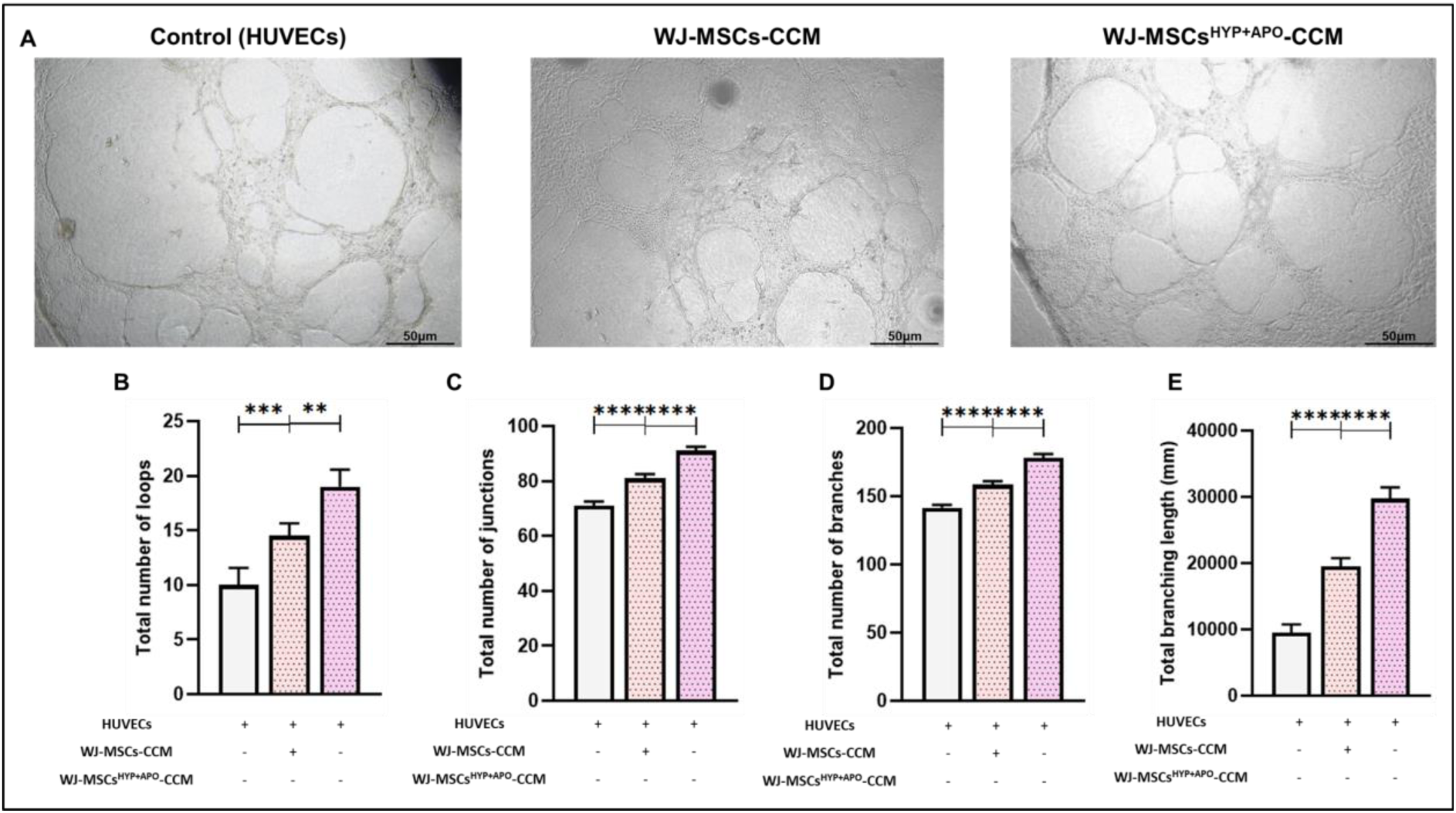
Tube formation assay of H_2_O_2_-treated HUVECs in the presence or absence of MSCs-CCM (WJ-MSCs-CCM, WJ-MSCS ^HYP+APO^-CCM). (A) Representative phase-contrast images of HUVECs cultured under different conditions: Control, WJ-MSCs-CCM, WJ-MSCs^HYP+APO^-CCM. The bar graphs represent the angiogenic parameters (B) total number of loops, (C) total number of junctions, (D) total number of branches, (E) total branching length (mm). Data shown represent the Mean±S.D of 5 independent experiments, with each experiment conducted in triplicate (technical replicates). Statistical analysis: Tukey’s multiple comparisons test; *≤0.05; **≤0.01; ***≤0.001; ****≤0.0001. Scale bar: 20X: 50μm. Abbreviations: BM: Bone marrow; WJ: Wharton’s Jelly; MSCs: Mesenchymal Stem Cells; HYP: Hypoxia; APO: Apoptosis; CCM: Culture Conditioned Media

**Figure S7:**
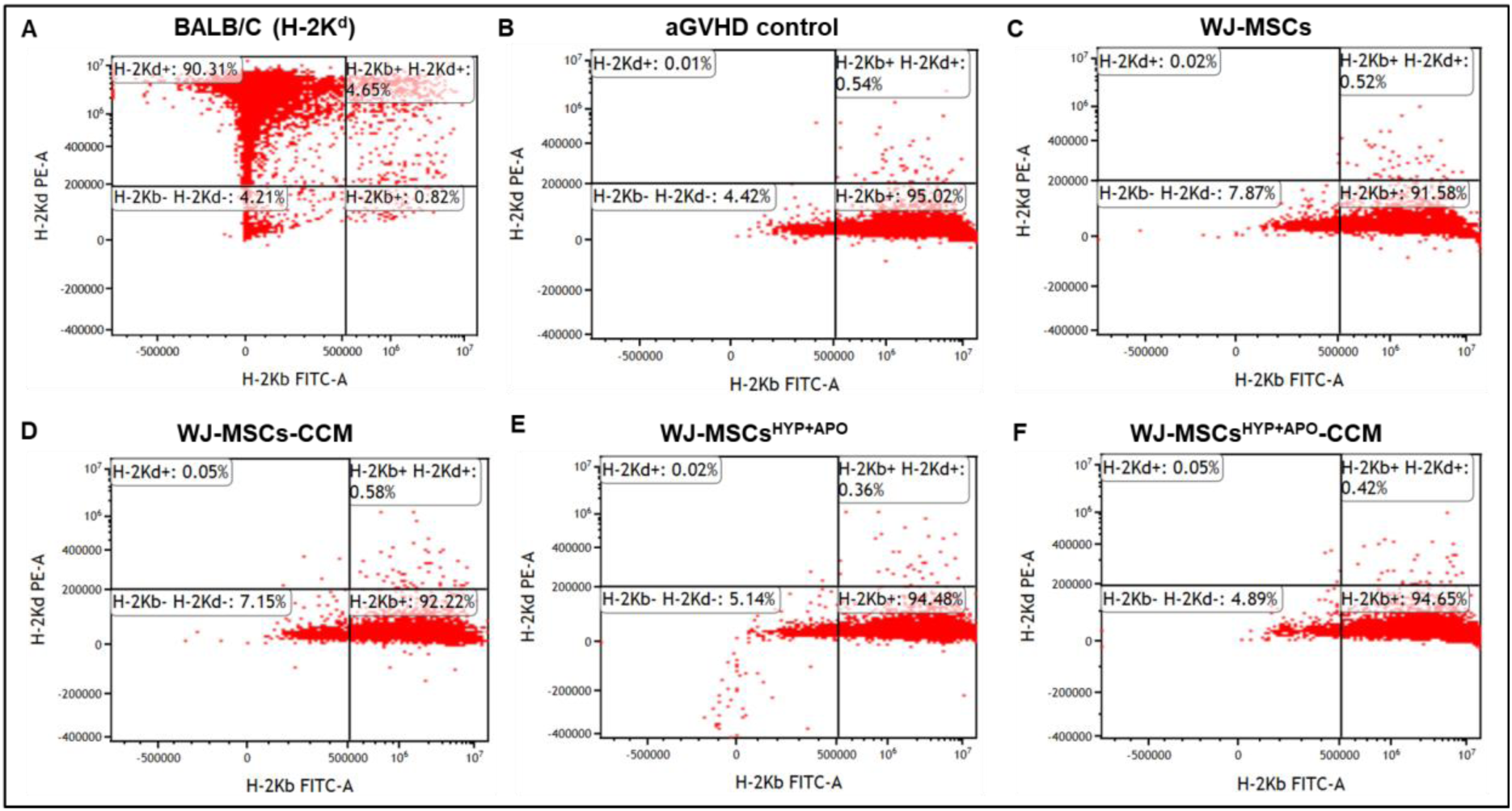
Effect of WJ-MSCs, WJ-MSCs^HYP+APO^, and their CCM on stem cell engraftment in a chemotherapy-based aGVHD murine model. The dot plots represent the chimerism in (A) BalB/c (without transplant). (B) aGVHD control. (C) WJ-MSCs. (D) WJ-MSCs-CCM. (E) WJ-MSCs ^HYP+APO^. (F) WJ-MSCs ^HYP+APO^-CCM. Abbreviations: WJ: Wharton’s Jelly; MSCs: Mesenchymal Stem Cells; HYP: Hypoxia; APO: Apoptosis; CCM: Culture-Conditioned Media

**Figure S8:**
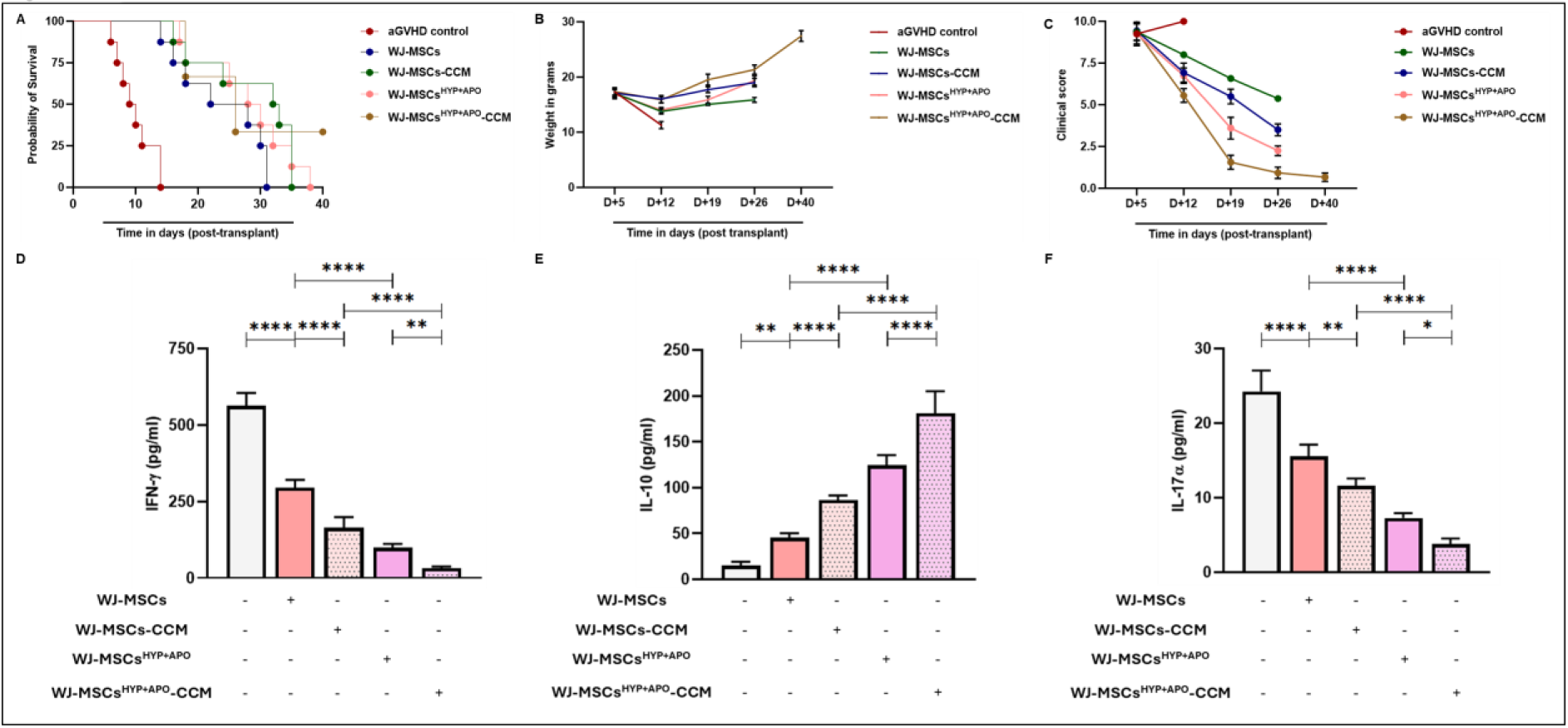
Effect of WJ-MSCs, WJ-MSCs^HYP+APO^, and their CCM in a chemotherapy-based aGVHD murine model. Line graphs represent (A) clinical score. (B) body weight. (C) overall survival. The bar graphs represent the concentration (pg/ml) of (D) IFN-γ. (F) IL-10. (E) IL-17α. Data presented as Mean ±S.D. (N= 8 mice /group). Abbreviations: WJ: Wharton’s Jelly; MSCs: Mesenchymal Stem Cells; HYP: Hypoxia; APO: Apoptosis; CCM: Culture-Conditioned Media

**Figure S9:**
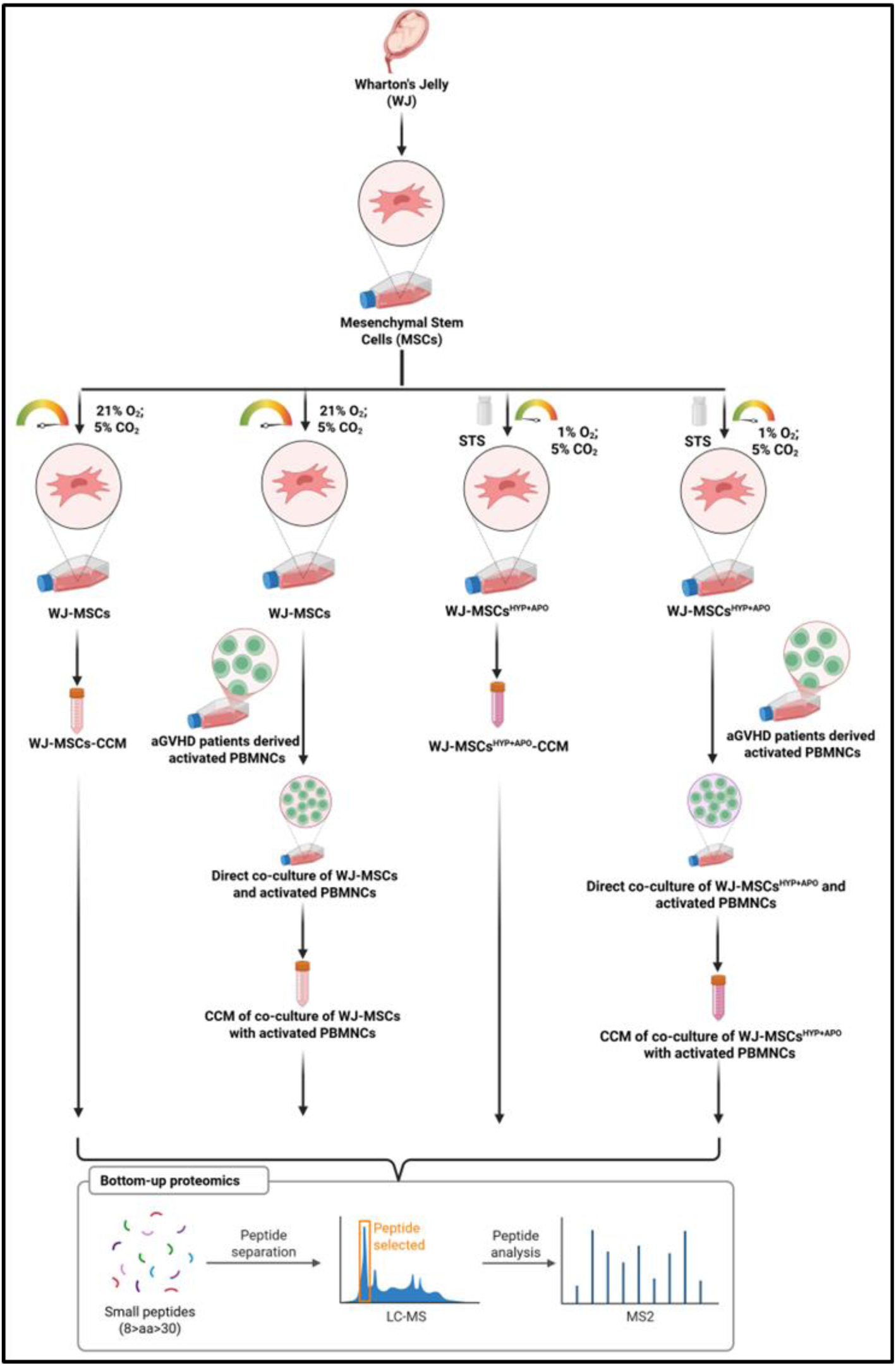
Schematic representation of the workflow adopted for proteomic profiling of CCM of WJ-MSCs, WJ-MSCs^HYP+APO^, and their co-culture with aGVHD patients-derived activated PBMNCs. Abbreviations: PBMNCs: Peripheral Blood Mononuclear Cells

**Figure S10:**
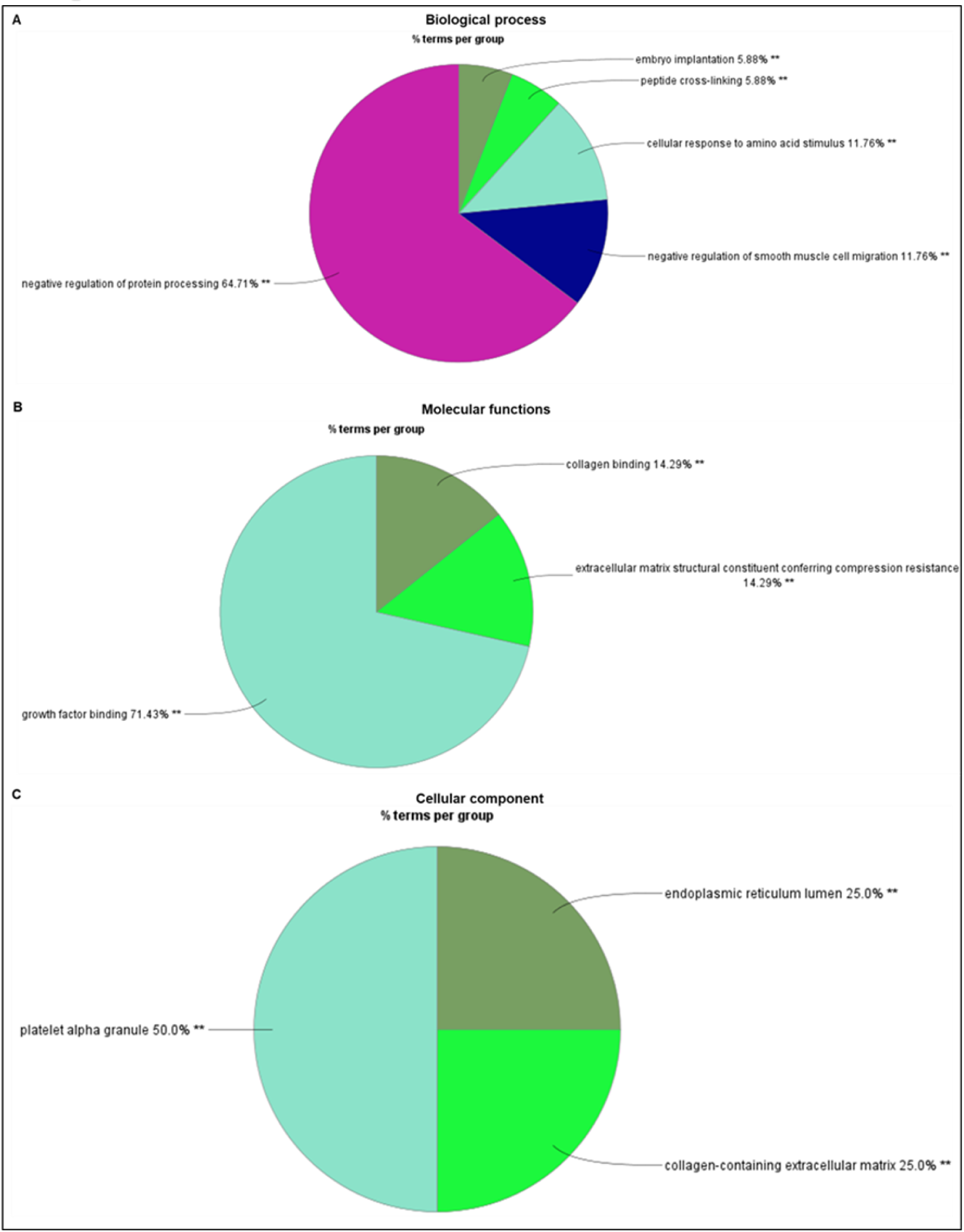
Gene Ontology (GO) analysis of upregulated proteins in WJ-MSCs-CCM compared to their WJ-MSCs^HYP+APO^-CCM. Pie charts depict. (A) Biological process. (B) Molecular function. (C) Cellular component. Data showed three independent experiments conducted with three different donors (biological replicates). **≤0.01

**Figure S11:**
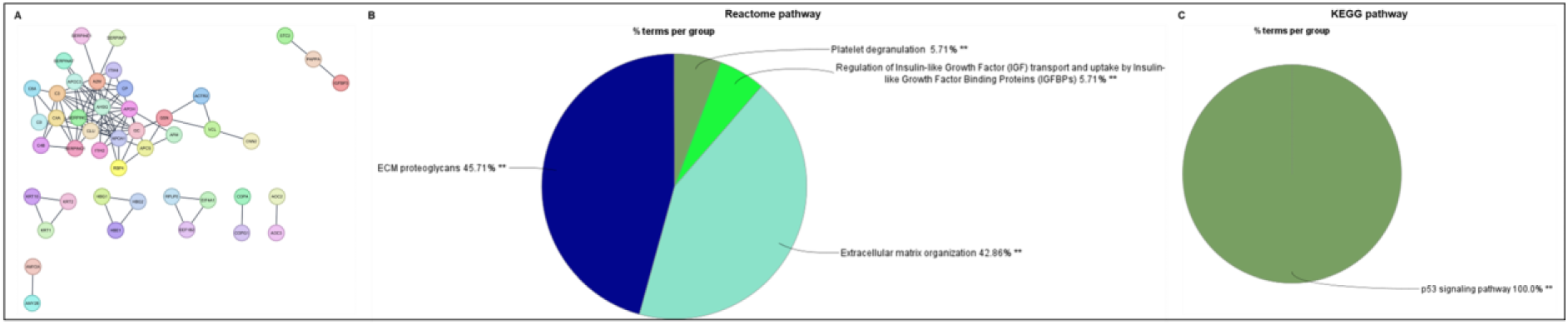
Functional enrichment analysis of upregulated proteins in WJ-MSCs-CCM compared to their WJ-MSCs^HYP+APO^-CCM. (A) STRING network. Pie charts depict (B) Reactome pathway. (C) KEGG pathway. (Data showed three independent experiments conducted with three different donors (biological replicates). **≤0.01

**Figure S12:**
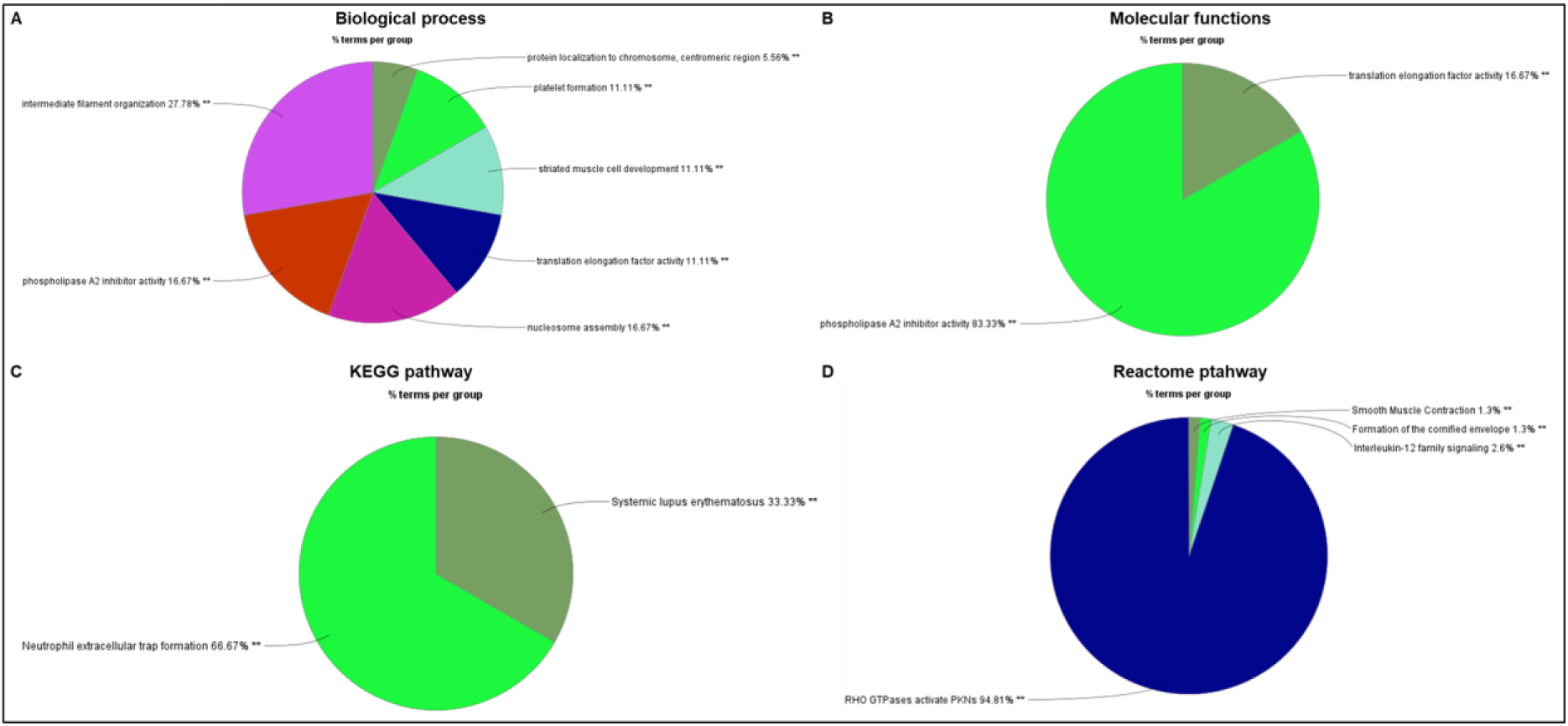
Gene Ontology (GO) and functional enrichment analysis of downregulated proteins in WJ-MSCs-CCM compared to WJ-MSCs^HYP+APO^-CCM. Pie charts depict (A) Biological process. (B) Molecular functions. (C) KEGG pathway. (D) Reactome pathway. Data showed three independent experiments conducted with three different donors (biological replicates).

**Figure S13:**
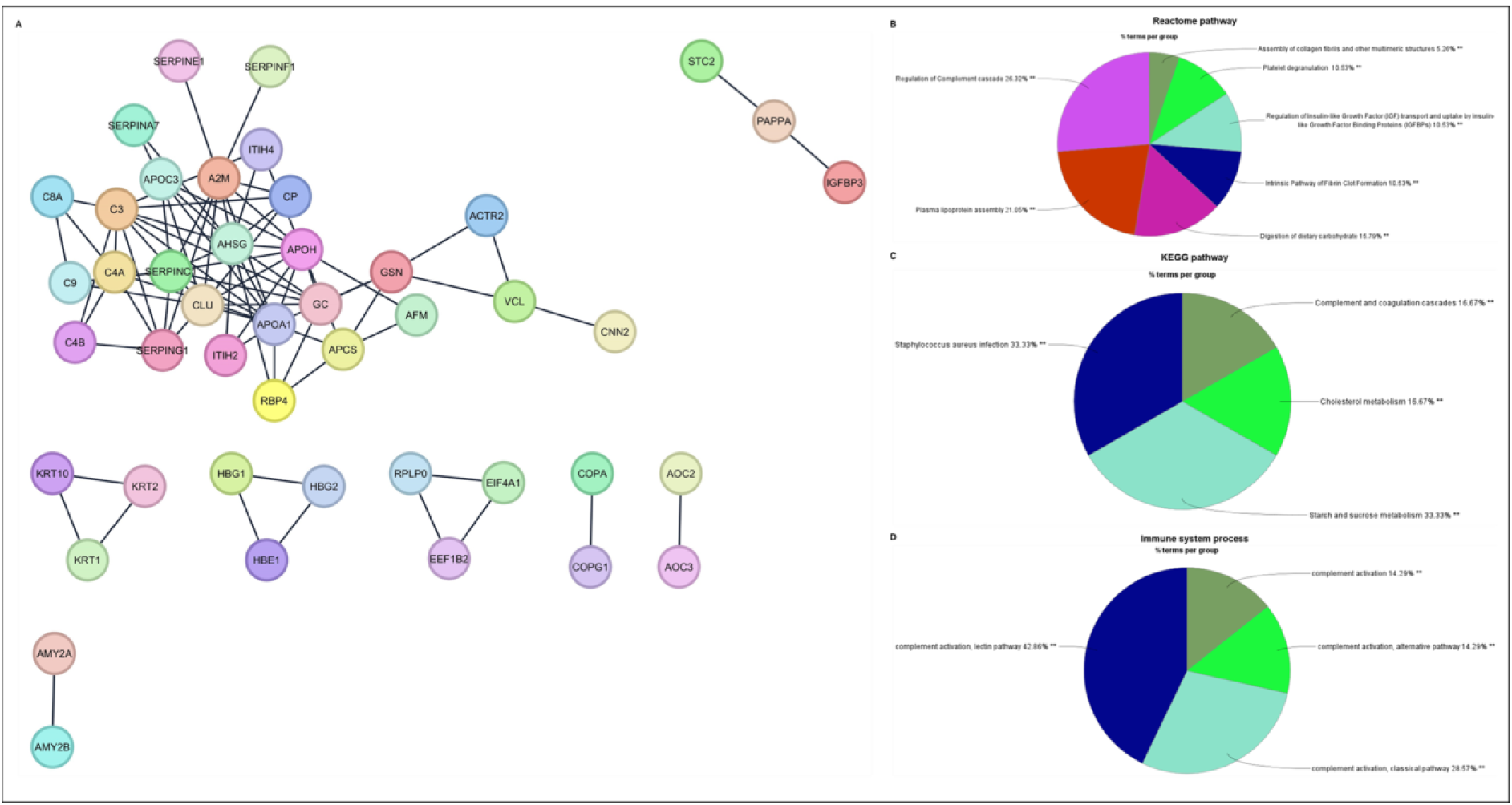
Functional enrichment analysis of upregulated proteins in CCM of coculture of WJ-MSCs with aPBMNCs compared to WJ-MSCs. (A) STRING network. Pie charts depict (B) the Reactome pathway. (C) KEGG pathway. (D) Immune system process. Data showed three independent experiments conducted with three different donors (biological replicates). **≤0.01

**Figure S14:**
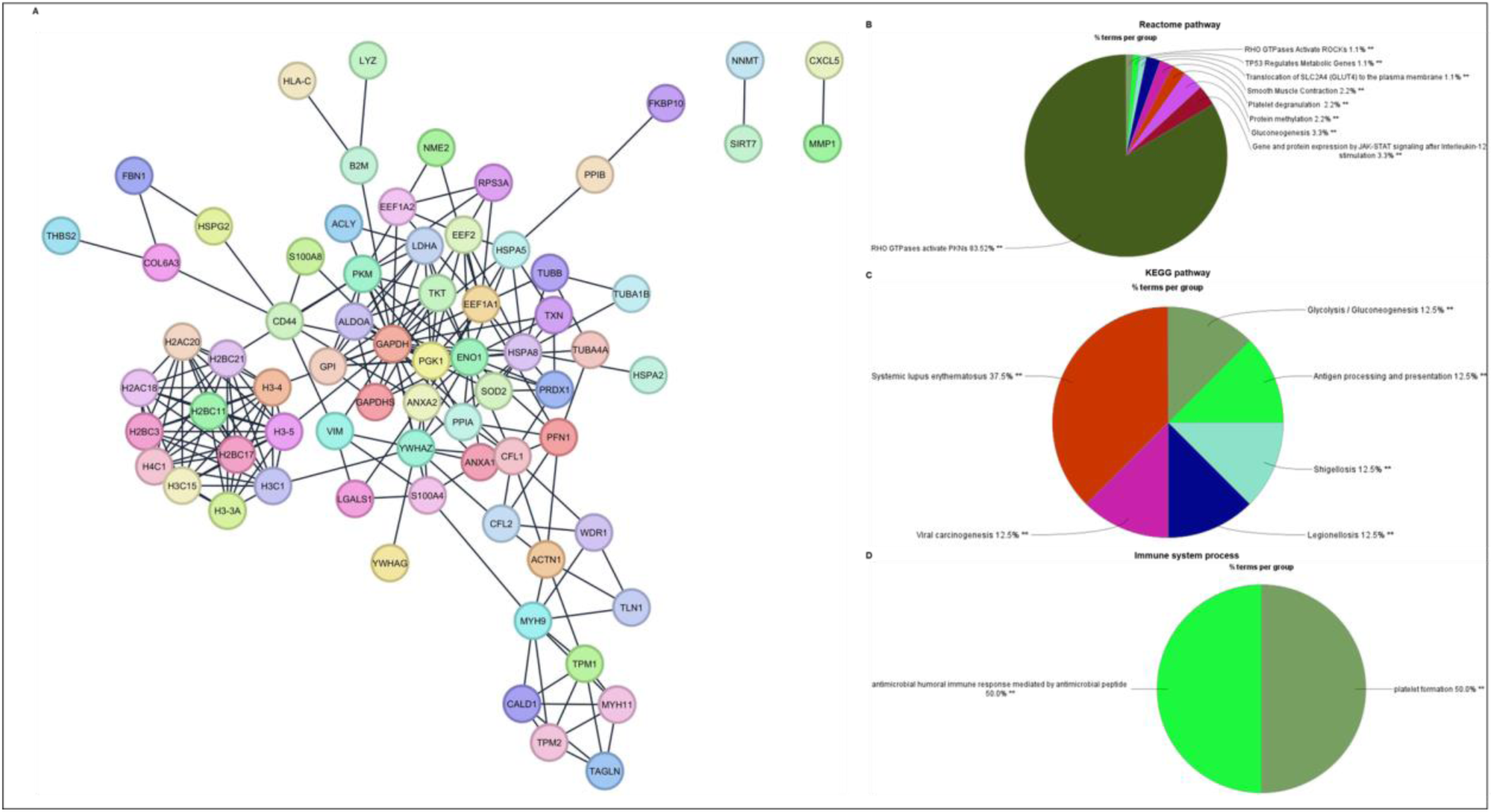
Functional enrichment analysis of downregulated proteins in the CCM of coculture of WJ-MSCs with aPBMNCs compared to WJ-MSCs. (A) STRING network. Pie charts depict (B) Reactome pathway. (C) KEGG pathway. (D) Immune system process. Data showed three independent experiments conducted with three different donors (biological replicates).

**Figure S15:**
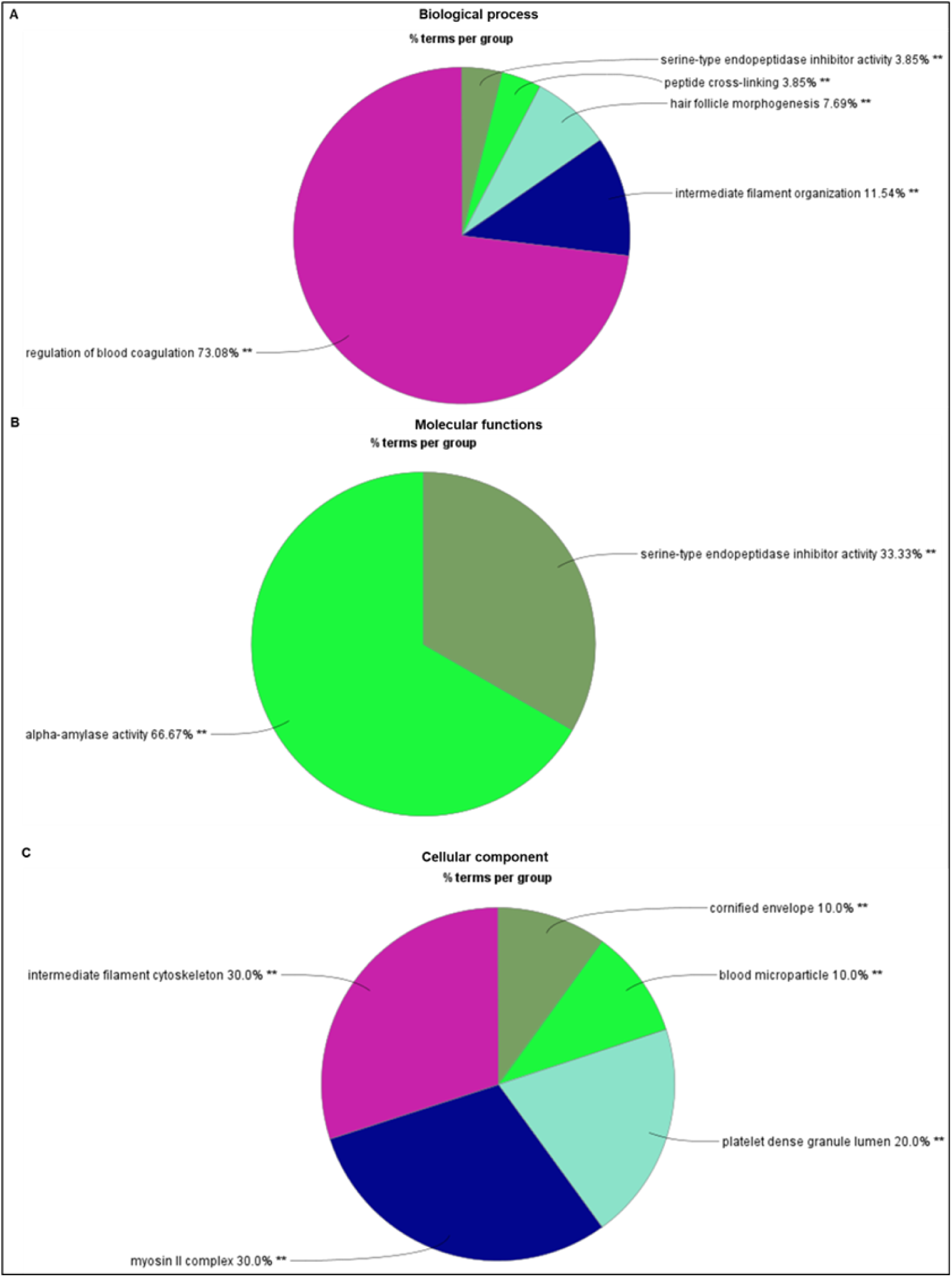
Gene Ontology (GO) analysis of upregulated proteins in WJ-MSCs^HYP+APO^-CCM compared to the CCM of coculture of WJ-MSCs^HYP+APO^ with aPBMNCs derived from aGVHD patients. Pie charts depict (A) Biological process. (B) Molecular function. (C) Cellular component. Data showed three independent experiments conducted with three different donors (biological replicates). **≤0.01

**Figure S16:**
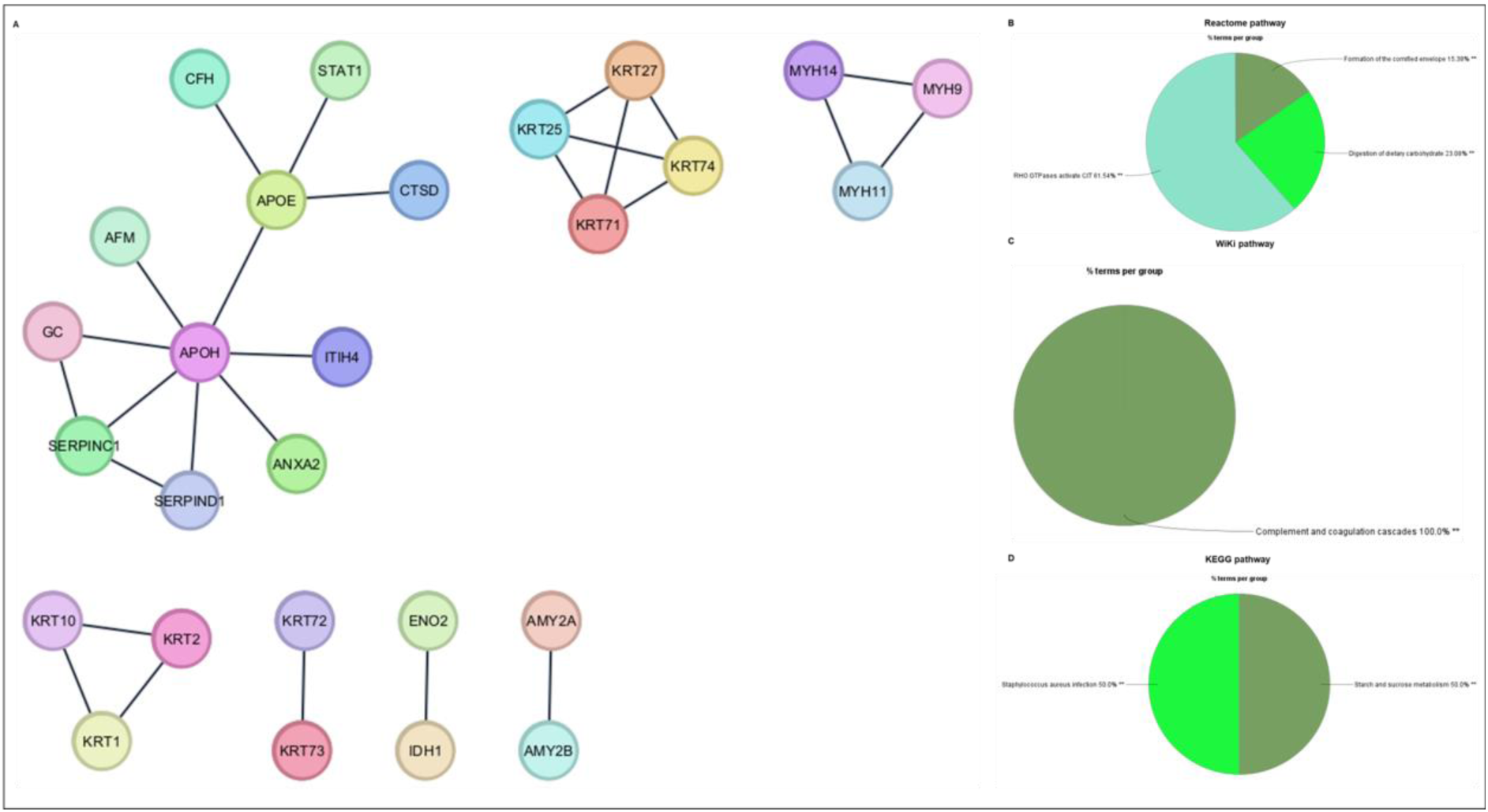
Functional enrichment analysis of upregulated proteins in WJ-MSCs^HYP+APO^-CCM compared to the CCM of coculture of WJ-MSCs^HYP+APO^ with aPBMNCs derived from aGVHD patients. (A) STRING network. Pie charts depict (B) Reactome pathway. (C) Wiki pathway. (D) KEGG pathway. Data showed three independent experiments conducted with three different donors (biological replicates). **≤0.01

**Figure S17:**
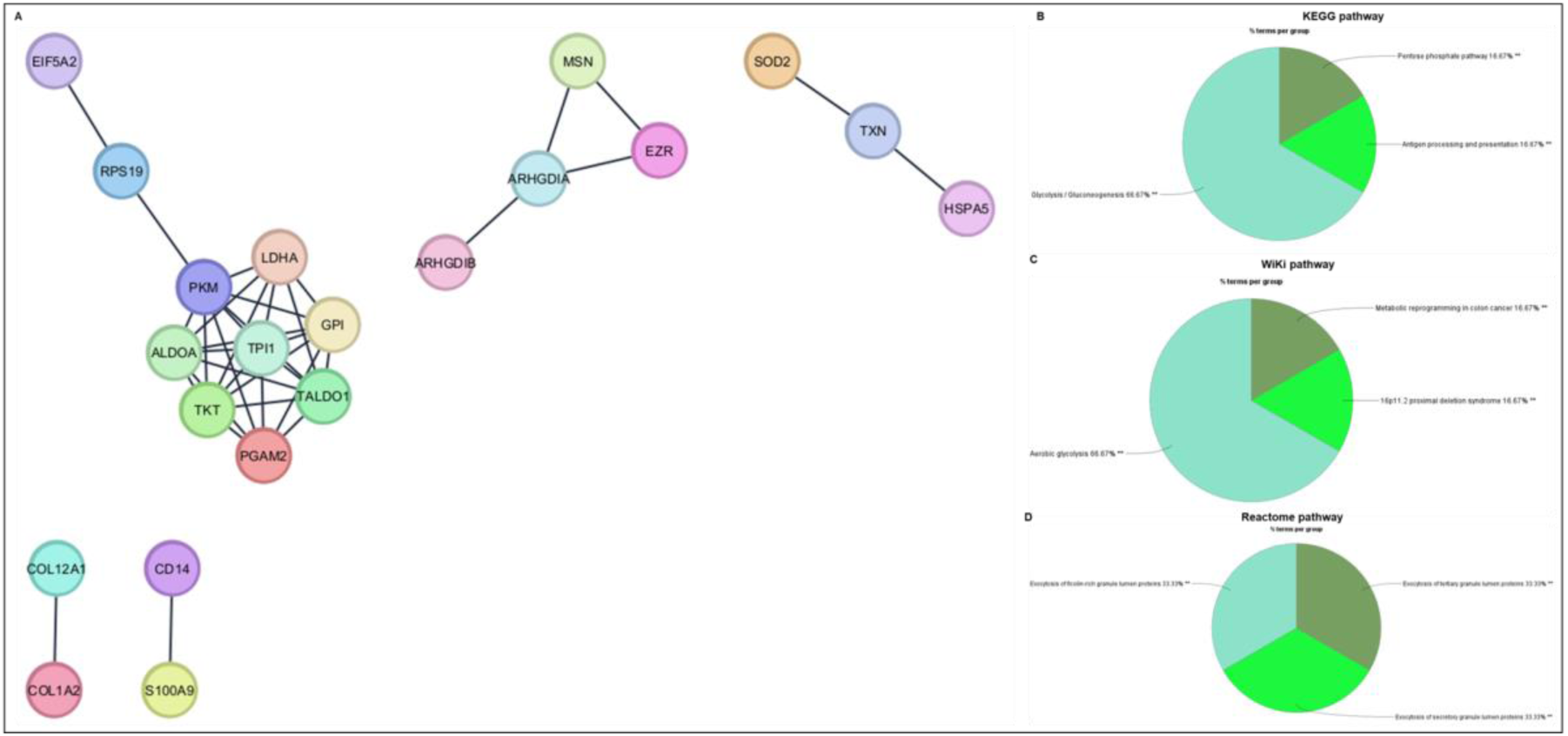
Functional enrichment analysis of downregulated proteins in WJ-MSCs^HYP+APO^-CCM compared to their CCM of coculture of WJ-MSCs^HYP+APO^ with aPBMNCs derived from aGVHD patients. (A) STRING network. Pie charts depict (B) KEGG pathway. (C) Wiki pathway. (D) Reactome pathway. Data showed three independent experiments conducted with three different donors (biological replicates). **≤0.01

**Figure S18:**
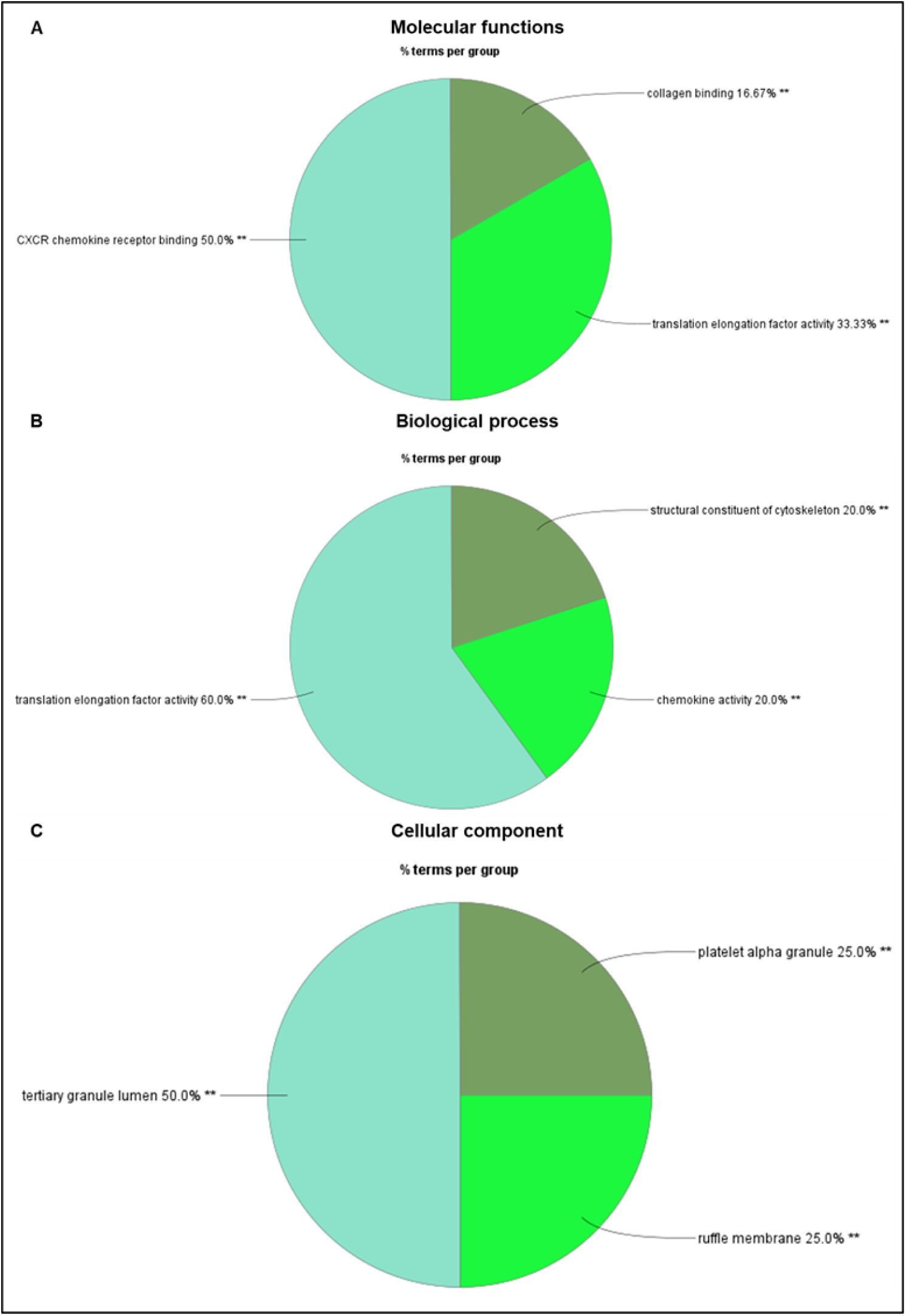
Gene Ontology (GO) analysis of upregulated proteins in the CCM of coculture of WJ-MSCs with aPBMNCs derived from aGVHD patients compared to the CCM of coculture of WJ-MSCs^HYP+APO^ with aPBMNCs derived from aGVHD patients. Pie charts depict (A) Molecular functions. (B) Biological process. (C) Cellular component. Data showed three independent experiments conducted with three different donors (biological replicates). **≤0.01

**Figure S19:**
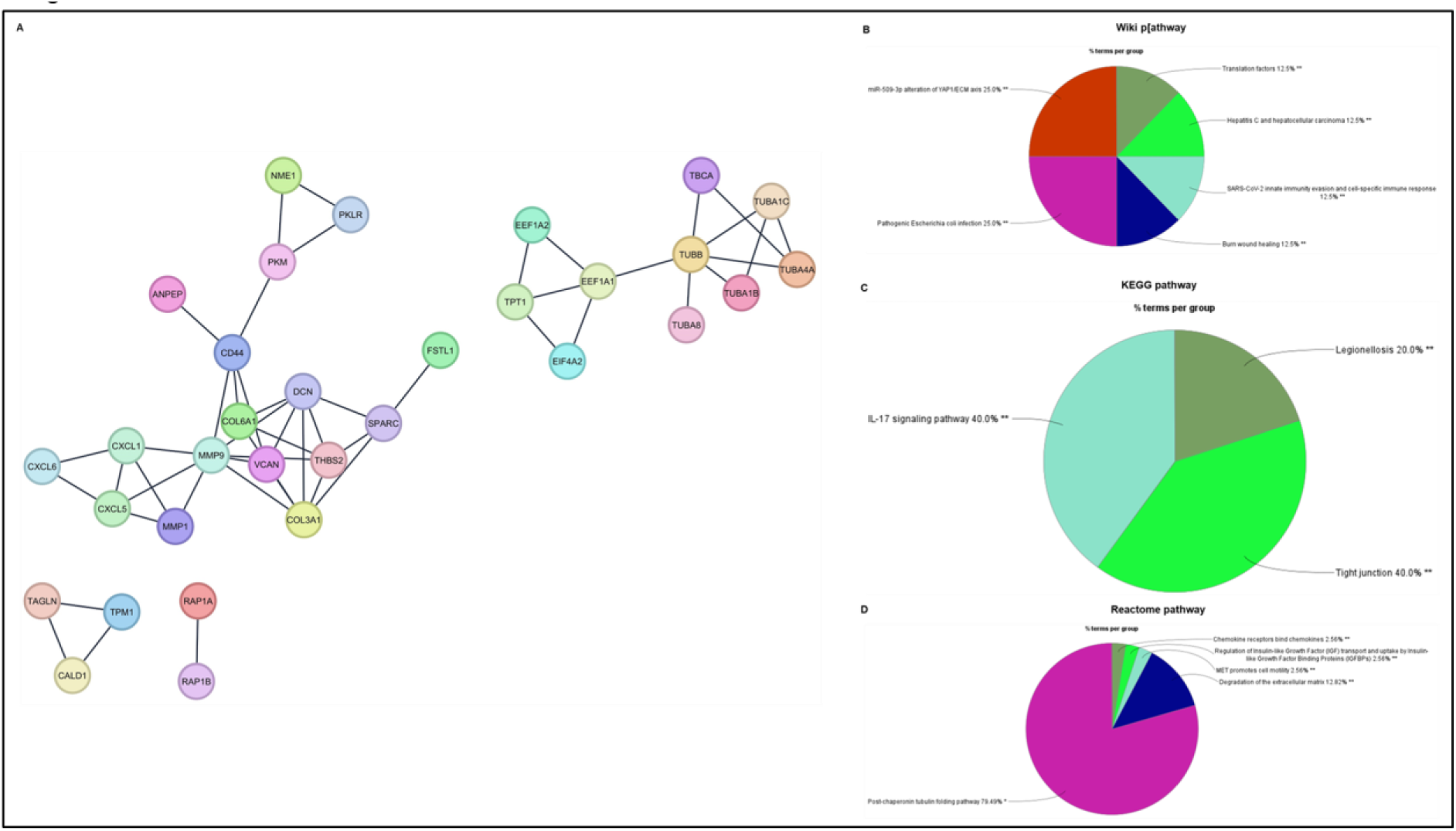
Functional enrichment analysis of upregulated proteins in the CCM of coculture of WJ-MSCs with aPBMNCs derived from aGVHD patients compared to the CCM of coculture of WJ-MSCs^HYP+APO^ with aPBMNCs derived from aGVHD patients. (A) STRING network. Pie charts depict (B) Wiki pathway. (C) KEGG pathway. (D) Reactome pathway. Data showed three independent experiments conducted with three different donors (biological replicates). **≤0.01

**Figure S20:**
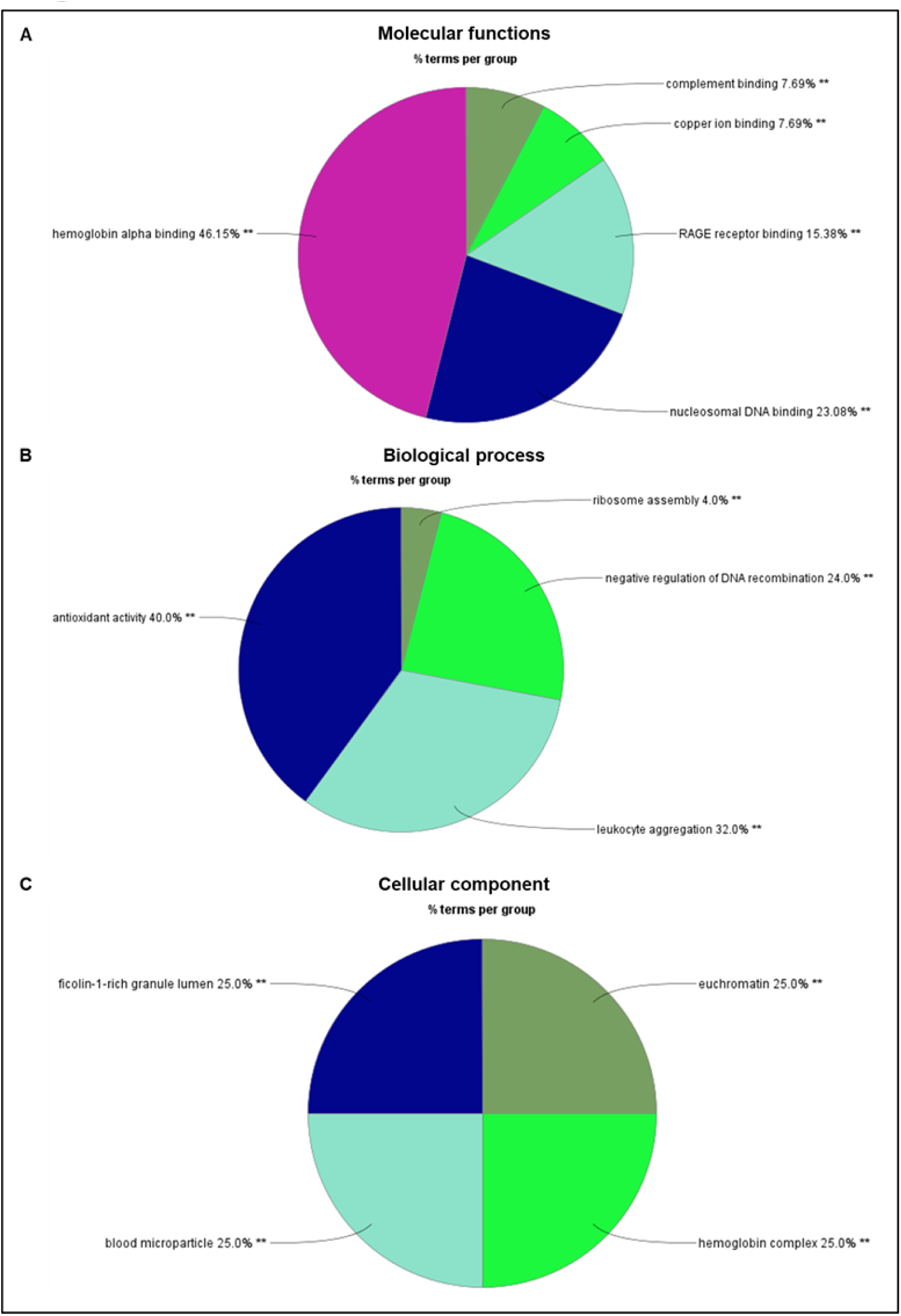
Gene Ontology (GO) analysis of downregulated proteins in the CCM of coculture of WJ-MSCs with aPBMNCs derived from aGVHD patients compared to the CCM of coculture of WJ-MSCs^HYP+APO^ with aPBMNCs derived from aGVHD patients. Pie charts depict (A) Molecular functions. (B) Biological process. (C) Cellular component. Data showed three independent experiments conducted with three different donors (biological replicates). **≤0.01

**Figure S21:**
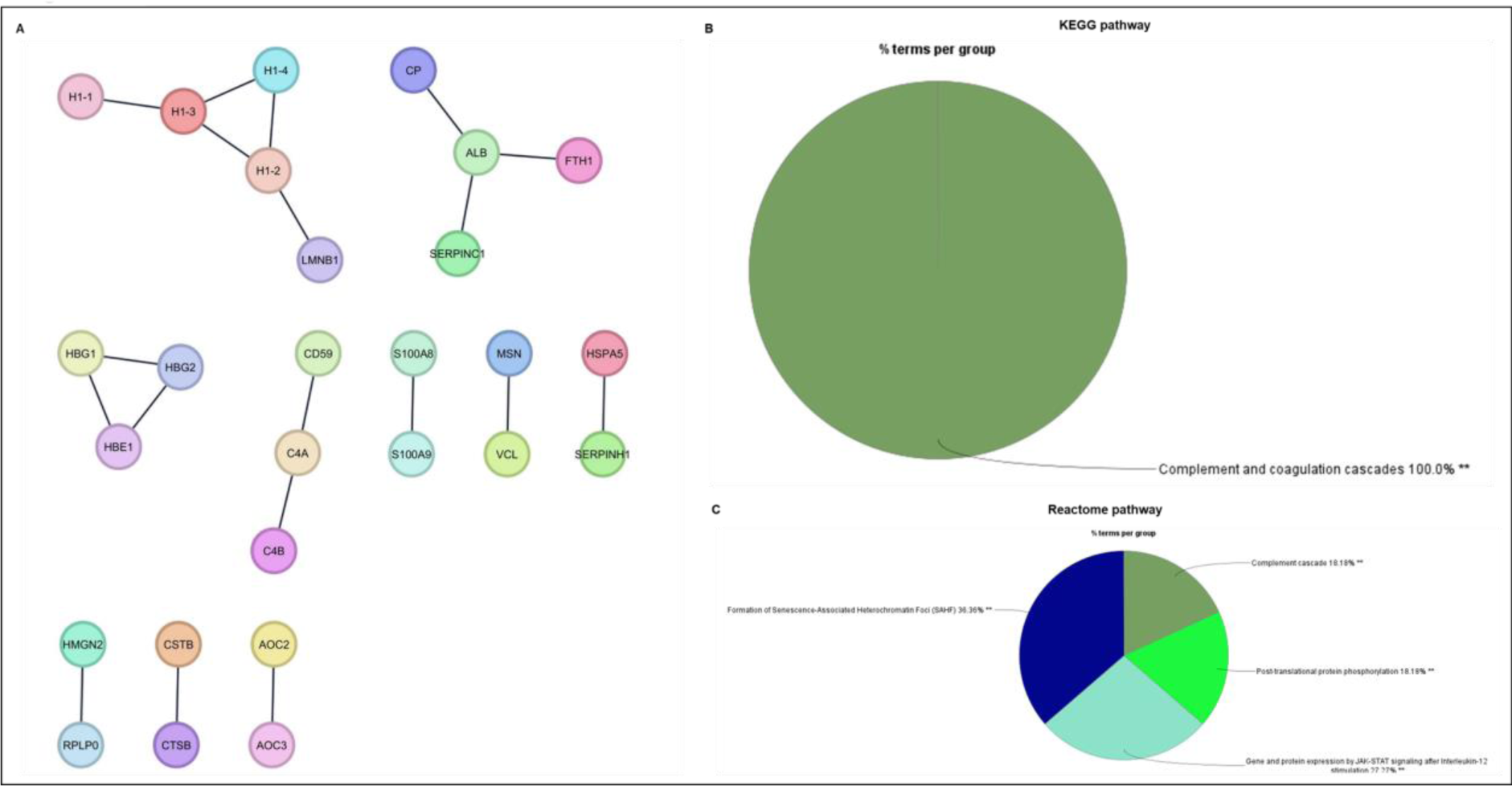
Functional enrichment analysis of downregulated proteins in the CCM of coculture of WJ-MSCs with aPBMNCs derived from aGVHD patients compared to the CCM of coculture of WJ-MSCs^HYP+APO^ with aPBMNCs derived from aGVHD patients. (A) STRING network. Pie charts depict (B) Wiki pathway. (C) KEGG pathway. (D) Reactome pathway. Data showed three independent experiments conducted with three different donors (biological replicates). **≤0.01

